# Inherited Causes of Clonal Hematopoiesis of Indeterminate Potential in TOPMed Whole Genomes

**DOI:** 10.1101/782748

**Authors:** Alexander G Bick, Joshua S Weinstock, Satish K Nandakumar, Charles P Fulco, Matthew J Leventhal, Erik L Bao, Joseph Nasser, Seyedeh M Zekavat, Mindy D Szeto, Cecelia Laurie, Margaret A Taub, Braxton D Mitchell, Kathleen C Barnes, Arden Moscati, Myriam Fornage, Susan Redline, Bruce M Psaty, Edwin K Silverman, Scott T Weiss, Nicholette D Palmer, Ramachandran S Vasan, Esteban G Burchard, Sharon LR Kardia, Jiang He, Robert C Kaplan, Nicholas L Smith, Donna K Arnett, David A Schwartz, Adolfo Correa, Mariza de Andrade, Xiuqing Guo, Barbara A Konkle, Brian Custer, Juan M Peralta, Hongsheng Gui, Deborah A Meyers, Stephen T McGarvey, Ida Yii-Der Chen, M. Benjamin Shoemaker, Patricia A Peyser, Jai G Broome, Stephanie M Gogarten, Fei Fei Wang, Quenna Wong, May E Montasser, Michelle Daya, Eimear E Kenny, Kari North, Lenore J Launer, Brian E Cade, Joshua C Bis, Michael H Cho, Jessica Lasky-Su, Donald W Bowden, L Adrienne Cupples, Angel CY Mak, Lewis C Becker, Jennifer A Smith, Tanika N Kelly, Stella Aslibekyan, Susan R Heckbert, Hemant K Tiwari, Ivana V Yang, John A Heit, Steven A Lubitz, Stephen S Rich, Jill M Johnsen, Joanne E Curran, Sally Wenzel, Daniel E Weeks, Dabeeru C Rao, Dawood Darbar, Jee-Young Moon, Russell P Tracy, Erin J Buth, Nicholas Rafaels, Ruth JF Loos, Lifang Hou, Jiwon Lee, Priyadarshini Kachroo, Barry I Freedman, Daniel Levy, Lawrence F Bielak, James E Hixson, James S Floyd, Eric A Whitsel, Patrick Ellinor, Marguerite R Irvin, Tasha E Fingerlin, Laura M Raffield, Sebastian M Armasu, Jerome I Rotter, Marsha M Wheeler, Ester C Sabino, John Blangero, L. Keoki Williams, Bruce D Levy, Wayne Huey-Herng Sheu, Dan M Roden, Eric Boerwinkle, JoAnn E Manson, Rasika A Mathias, Pinkal Desai, Kent D Taylor, Andrew Johnson, Paul L Auer, Charles Kooperberg, Cathy C Laurie, Thomas W Blackwell, Albert V Smith, Hongyu Zhao, Ethan Lange, Leslie Lange, James G Wilson, Eric S Lander, Jesse M Engreitz, Benjamin L Ebert, Alexander P Reiner, Vijay G Sankaran, Sidd Jaiswal, Goncalo Abecasis, Pradeep Natarajan, Sekar Kathiresan, on behalf of the NHLBI Trans-Omics for Precision Medicine (TOPMed) Consortium

## Abstract

Age is the dominant risk factor for most chronic human diseases; yet the mechanisms by which aging confers this risk are largely unknown.^1^ Recently, the age-related acquisition of somatic mutations in regenerating hematopoietic stem cell populations was associated with both hematologic cancer incidence^2–4^ and coronary heart disease prevalence.^5^ Somatic mutations with leukemogenic potential may confer selective cellular advantages leading to clonal expansion, a phenomenon termed ‘Clonal Hematopoiesis of Indeterminate Potential’ (CHIP).^6^ Simultaneous germline and somatic whole genome sequence analysis now provides the opportunity to identify root causes of CHIP. Here, we analyze high-coverage whole genome sequences from 97,691 participants of diverse ancestries in the NHLBI TOPMed program and identify 4,229 individuals with CHIP. We identify associations with blood cell, lipid, and inflammatory traits specific to different CHIP genes. Association of a genome-wide set of germline genetic variants identified three genetic loci associated with CHIP status, including one locus at *TET2* that was African ancestry specific. *In silico*-informed *in vitro* evaluation of the *TET2* germline locus identified a causal variant that disrupts a *TET2* distal enhancer. Aggregates of rare germline loss-of-function variants in *CHEK2*, a DNA damage repair gene, predisposed to CHIP acquisition. Overall, we observe that germline genetic variation altering hematopoietic stem cell function and the fidelity of DNA-damage repair increase the likelihood of somatic mutations leading to CHIP.

---

The U.S. National Heart, Lung, and Blood Institute (NHLBI) Trans-omics for Precision Medicine (TOPMed) project seeks to use high-coverage (>35x) whole genome sequencing (WGS) and molecular profiling to improve fundamental understanding of heart, lung, blood, and sleep disorders.^7^ Within the TOPMed program, we designed a study to detect CHIP from blood DNA-derived WGS in 97,691 individuals across 52 largely observational epidemiologic studies to discover the inherited genetic causes and phenotypic consequences of CHIP (**Supplementary Table 1**).

To confidently identify somatic mutations in blood-derived DNA, we analyzed TOPMed WGS data with the GATK MuTECT2 somatic variant caller.^8^ We performed several quality control steps to identify and remove sequencing artifacts and germline mutations from the call set (see **Methods**). We used previously described methods to identify CHIP carriers on the basis of a pre-specified list of leukemogenic driver mutations in genes known to promote clonal expansion of hematopoietic stem cells (**Supplementary Table 2**).^5^

In total, we identified 4,938 CHIP mutations in 4,229 individuals (**Supplementary Table 3**). Consistent with prior reports, >75% of these CHIP mutations were in one of three genes, *DNMT3A, TET2*, and *ASXL1*. Approximately 15% of these CHIP mutations were in the five next most frequent genes (*PPM1D, JAK2, SF3B1, SRSF2* and *TP53*, **Figure 1a**). Amongst these 8 most commonly mutated genes, there was marked heterogeneity in clonal fraction. For example, among the top three genes, *DNMT3A* CHIP clonal fraction of the peripheral blood was ∼25% smaller (p= 1.3 × 10^−15^) and *TET2* clones were ∼14% smaller (p=2.1 × 10^−4^) than *ASXL1*, implicating the presence of driver mutation gene-specific differences in clonal selection (**Figure 1b**). Amongst the 4,229 individuals with CHIP driver mutations, 3,822 (90%) had a single mutation identified, and only 54 individuals (1%) had 3 or more CHIP driver mutations (**Figure 1c**).

**Fig. 1.**
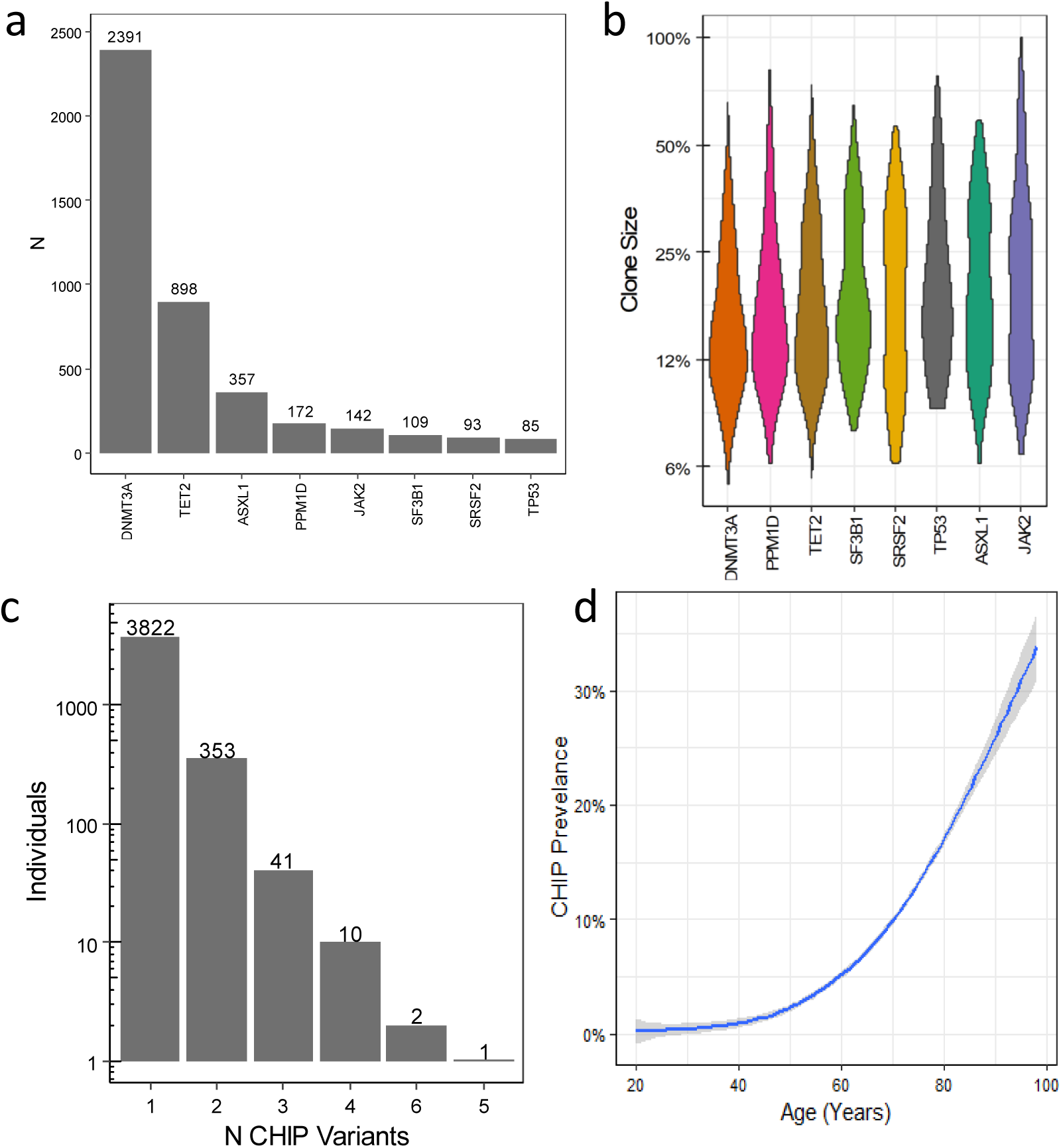
Characterizing CHIP in TOPMed Genomes. **a**, 97,631 peripheral blood samples were whole genome sequenced and called with a somatic variant caller. CHIP was identified through the curation of CHIP driver variants. Driver variants in DNMT3A and TET comprised more than half of the CHIP calls. **b**, There was marked heterogeneity of CHIP clone size as measured by allele fraction (y axis) by CHIP driver gene. **c**, 90% of individuals with CHIP had only one CHIP driver mutation identified. d, CHIP prevalence increased with age (p<10^−300^).

CHIP prevalence was strongly correlated with age at blood draw (p < 10^−300^, **Figure 1d**). The CHIP-age association was highly consistent across studies (**Extended Data Figure 1**) and comparable to previous reports^2–4^ which detected CHIP by whole exome sequencing (**Extended Data Figure 2**). Consistent with prior studies, history of smoking was associated with increased CHIP odds (OR = 1.18, p=5 × 10^−5^) whereas Hispanic ancestry and East Asian ancestry were each associated with reduced CHIP odds (OR = 0.50, p=0.008 and OR = 0.56, p=0.001 respectively). Male sex was associated with increased CHIP in univariate models, but this association did not persist in multi-variate models (**Supplementary Table 4**)

We considered whether there were differences in the CHIP age distributions by mutational mechanisms and driver gene. Carriers of frameshift CHIP mutations were on average older individuals than carriers of single nucleotide CHIP mutations (Wilcox rank sum test: p=0.01). *JAK2* CHIP carriers were the youngest. Relative to *JAK2, ASXL1* and *TET2* carriers were 3.3 and 3.9 years older respectively (p=0.01, p=9.1 × 10^−^ 4), while *PPM1D, SF3B1* and *SRSF2* carriers were 5.0, 6.9 and 7.7 years older (p=5.7 × 10^−4^, 1.8 × 10^−6^, 1.3 × 10^−4^) (**Extended Data Figure 3**).

CHIP is typically distinguished from other clonal hematologic disorders based on the absence of cytopenia, dysplasia, and neoplasia.^6^ We considered whether there were sub-clinical hematologic correlates of CHIP that might have utility in identifying CHIP carriers. We observed a modest increase in total white blood cell count (p=1.1 × 10^−5^) and a modest decrease in hemoglobin (p=0.04), among those with CHIP compared to those without (**Figure 2a, Supplemental Table 5**). In aggregate, CHIP driver mutations were associated with increased red blood cell distribution width (RDW, p=3.0 × 10^−5^) consistent with prior observations.^9^ Notably, RDW is a hematologic parameter that increases with age and predicts overall mortality and poor clinical outcomes in the setting of CVD and in older adults.^10^

**Fig. 2.**
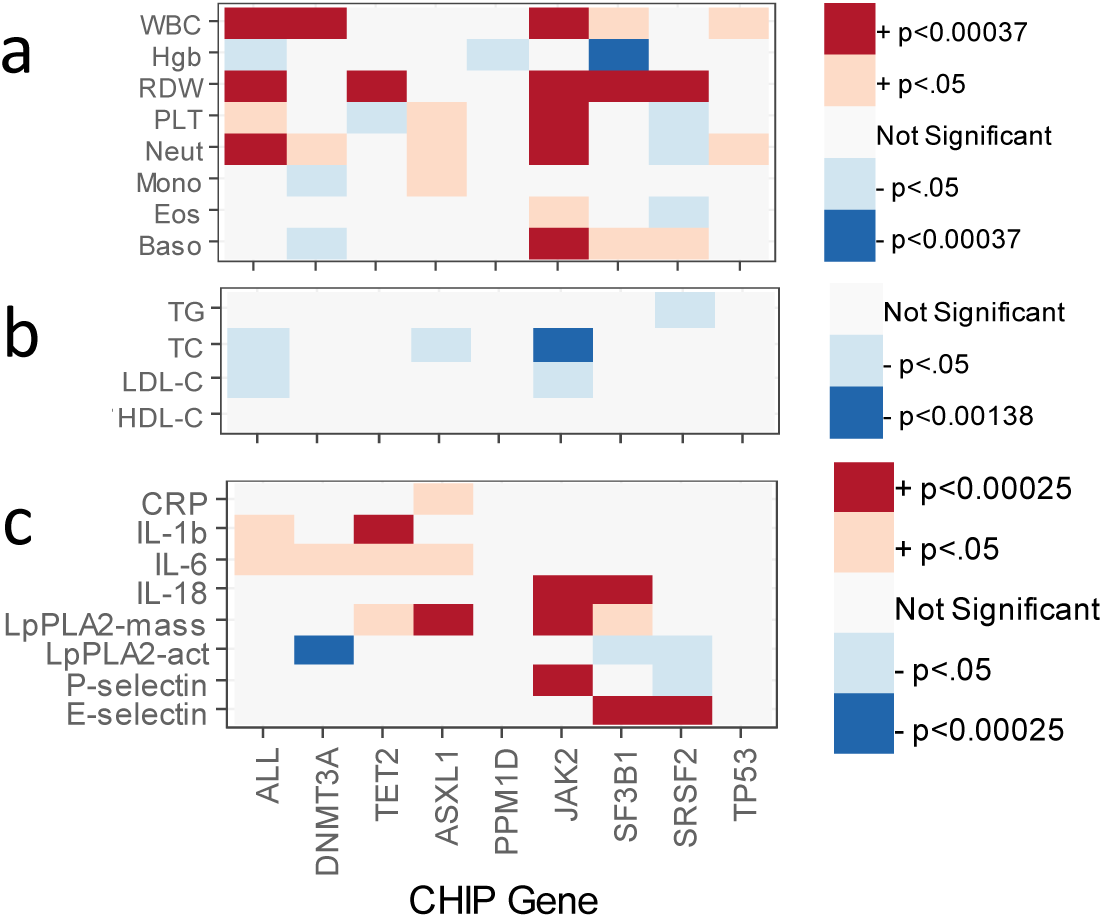
CHIP associates with Blood, Lipid, and Inflammatory traits. **a**, CHIP consistently associated with increased Red Cell Distribution Width (RDW). *JAK2, SF3B1* and *SRSF2* showed driver gene specific effects on blood traits. **b**, CHIP status was not consistently associated with lipid traits, other than *JAK2* CHIP which was associated with decreased total cholesterol and a trend towards decreased LDL and Triglycerides. **c**, CHIP status is associated with inflammatory markers, however notable heterogeneity existed across CHIP mutations.

Disaggregating by the 8 most common driver genes, we noted consistently elevated blood cell counts and specifically associations with increased platelets (p=2.5 × 10^−77^), basophils (p= 6.0 × 10^−18^) and neutrophils (p=8.7 × 10^−9^) in *JAK2* CHIP mutation carriers.

Given the prior association of CHIP with atherosclerotic cardiovascular disease^5^, we asked whether CHIP carriers had altered lipid profiles. Consistent with prior reports^5^, we observed negative correlations between *JAK2* and total cholesterol (p=5.1 x10^−4^) and LDL (p=0.0014) but no other significant associations (**Figure 2a, Supplemental Table 6**).

Hypercholesterolemic mice receiving *Tet2*-/- bone marrow have marked upregulation of macrophage cytokine signaling compared to hypercholesterolemic mice receiving wild-type bone marrow and pharmacologic manipulation of the IL-6/IL-1b axis mediates atherogenesis in mice.^5,11^ As a result, we sought to define the human inflammatory profile of CHIP carriers (**Figure 2a, Supplemental Table 7**). In aggregate, CHIP was associated with increased IL-6 (p=0.0035) and nominally associated with increased IL-1b (p=0.026). There was no association of CHIP with quantitative C-reactive protein (CRP) levels and elevated CRP did not reliably identify carriers of CHIP (AUC: 0.55; for cutoff of CRP>2: PPV=6.3%, sensitivity=60%). CHIP driver gene-specific analyses highlighted notable differences. For example, *TET2* CHIP carriers were associated with significantly increased IL-1b (p=2.4 × 10^−4^). Carriers of *JAK2* CHIP and *SF3B1* had increased circulating IL-18 (p=1.3×10^−4^ and 1.27 ×10^−20^ respectively).

Germline genetic variants have been previously associated with clonal hematopoiesis, defined either by somatic mosaicism of SNVs and indels^12^ or by chromosomal rearrangements with appreciable clonal fraction^13^, in individuals of European ancestry, and identified variants at a single locus, *TERT*, that associates with clonal hematopoiesis. Given the distinct association of clonal hematopoiesis with known leukemogenic mutations (i.e., CHIP) with both cancer^14^ and atherosclerotic cardiovascular disease^5^, we sought to specifically discover germline genetic variations conferring increased risk for CHIP acquisition. We performed a single variant genome-wide association analysis in a subset of 65,405 individuals (3,831 CHIP driver cases) where the likelihood of having a CHIP mutation was >1% (see **Methods**). The trait heritability explained by the analysis with LD score-regression was 3.6%.

Our WGS-based association analysis of CHIP replicated the lead variant of the single locus previously associated at genome wide significance with clonal hematopoiesis (defined based on somatic mosaicism of SNVs and indels),^12^ rs34002450 (OR 1.2, p=2.0 × 10^−13^). rs34002450 is in strong LD (r^2^=0.55) with our lead variant at this locus rs7705526, a common variant (MAF 0.29) in the 5^th^ intron of *TERT*, which encodes telomere enzyme reverse transcriptase. In TOPMed, carriers of the rs34002450 A (minor) allele have a 1.3-fold risk of developing CHIP (p=8.4x10^−24^). This variant was previously significantly associated with increased leukocyte telomere length.^15^ This variant was also associated with myeloproliferative neoplasms (MPN, see Bao et al, co-submitted manuscript) and clonal chromosomal mosaicism^13^. In a phenome-wide association analysis (PheWAS) of rs34002450 in UK Biobank, we identified significant increased risk of MPN (p=2.6 × 10^−13^), uterine leiomyoma (p=3.2 × 10^−9^), brain cancer (p=3.6 × 10^−8^) and a decreased risk of Seborrheic keratosis (p=1.4 × 10^−7^).

We performed a conditional analysis of the 14 other genome-wide significant SNPs at the TERT locus, conditioning on the lead SNP, to see if there were any additional signals that were independent of rs7705526. We identified a second intronic TERT variant rs13167280 (MAF 0.11, r^2^=0.2 with rs7705526) that independently associates with CHIP status (OR 1.3, p=6.1×10^−10^; conditional OR: 1.1, p=4.7 × 10^−4^).

In the TOPMed single-variant association analysis, we additionally identified 2 other novel genome-wide significant genetic loci, including one locus on chromosome 3 in an intergenic region spanning *KPNA4*/*TRIM59* and one locus on chromosome 4 near *TET2* (**Figure 3, Extended Data figure 4, Supplementary Table 8**).

**Fig. 3.**
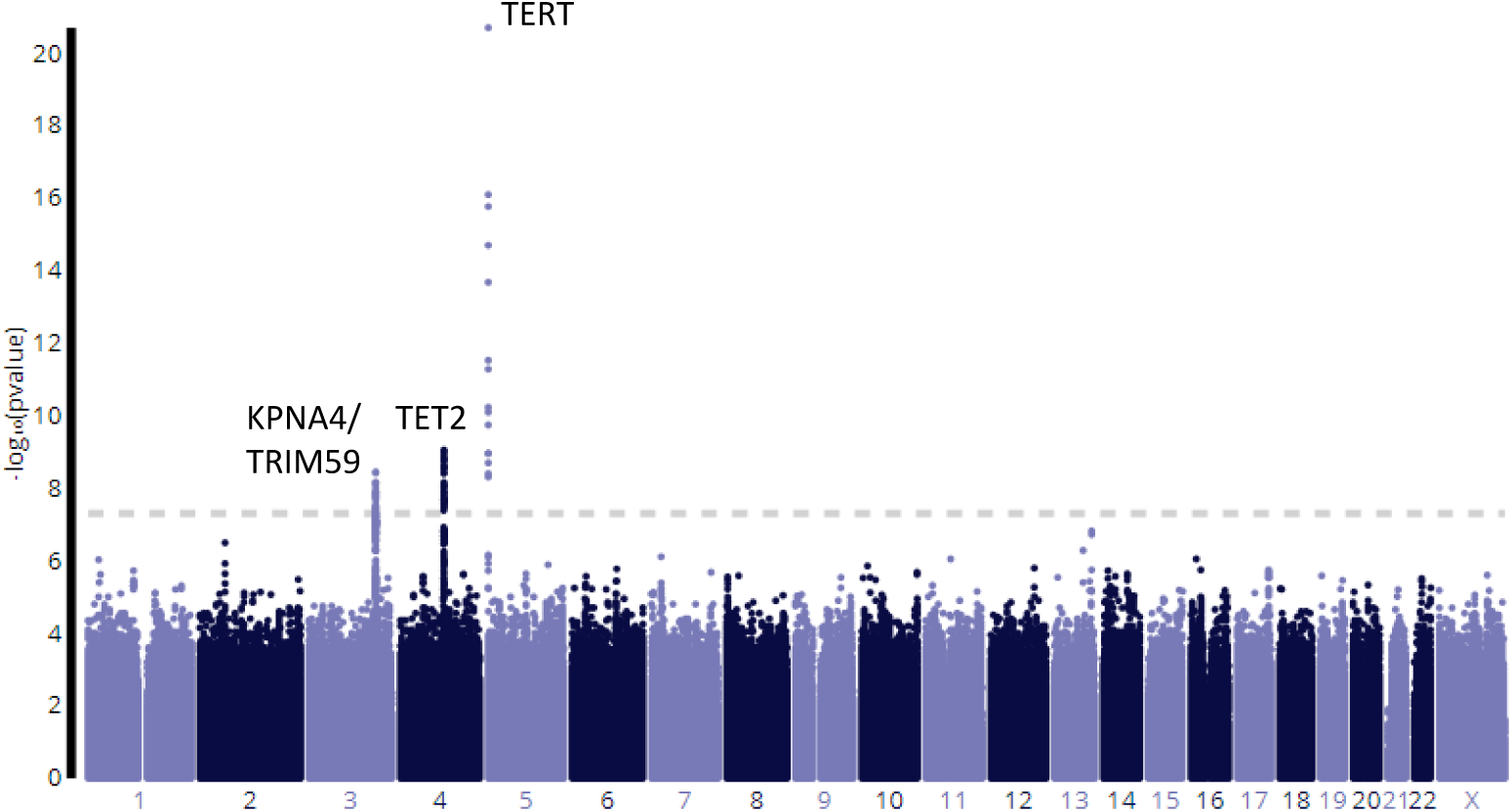
Genetic Determinants of CHIP. Single variant genetic association analyses of CHIP identified three genome wide significant loci.

rs1210060191 is a common variant (MAF 0.54) in a locus with an association signal that spans a 300kb region that includes *KPNA4, TRIM59, IFT80, and SMC4.*The lead variant is a 1 bp intronic deletion in *TRIM59*. Carriers of the del(T) allele have a 1.16-fold increased risk of CHIP (p=5.3×10^−10^) Variants in LD with this variant have been identified as associated with MPN (Bao et al, co-submitted manuscript) No other significant phenotype associations were noted in UK Biobank PheWAS analyses.

rs144418061 is an African ancestry specific variant (MAF 0.035 in African Ancestry samples, not present in non-African-ancestry samplies) in an intergenic region near *TET2.* Carriers of the A allele, have a 2.4-fold increased risk for CHIP (p=4.0×10^−9^). The association is equally robust for *DNMT3A* CHIP, *TET2* CHIP and *ASXL1* CHIP, suggesting that the germline variant does not specifically predispose to *TET2* CHIP. Although other variants in the vicinity of *TET2* have been associated with MPN (Bao et al, co-submitted manuscript), this variant has not been previously identified as associated with any traits in the literature.

We considered whether there might be germline variants that predispose to specific CHIP driver mutations by performing a GWAS on *DNMT3A* and *TET2* CHIP. We identified a single novel locus for *DNMT3A* chip at rs2887399 in an intron of T-cell leukemia/lymphoma 1A (TCL1A). Carriers of the T allele (MAF 0.26) are at 1.23 fold increased risk of acquiring a *DNMT3A* CHIP mutation (p=3.9 × 10^−9^). Intriguingly carriers of the T allele are at decreased risk of acquiring a *TET2* CHIP mutation (OR: 0.82, p=.0012), and consequently it was not identified in the primary CHIP GWAS analysis. This variant has also recently been associated with mosaic loss of chromosome Y.^16^

As single-variant analyses have limited power to detect rare-variant associations, we next performed several types of variant aggregation association tests. We performed a transcriptome-wide association analysis to quantify the relationship between changes in gene expression and genetic predisposition to CHIP.^17,18^ This approach identified the Chr3 *KPNA4/TRIM59* locus and six additional loci including: *AHRR, ASL, KREMN2, LEAP2, JSRP1, RASEF*. (**Extended Data Fig. 5-6)** *AHRR* directs hematopoietic progenitor cell expansion and differentiation.^19^

We also performed gene-based association tests for aggregations of rare (MAF<0.1%) putative loss-of-function (pLOF) germline variants within genes for CHIP presence. We considered all genes where at least ten individuals in the dataset were pLOF carriers. (15,031 genes with at least 10 LoF carriers, alpha=3.3 × 10^−6^) We filtered out all variants that were putatively somatic from the germline call set. Although no genes reached exome-wide significance, the top associated gene was DNA damage repair gene *CHEK2* (OR 1.7, p=1.3×10^−5^, **Supplementary Table 9**). Rare germline variants in *CHEK2* are implicated in the pathogenesis of a diverse set of hematologic and solid tumor malignancies.^17,18^ Common variants in *CHEK2* have previously been associated with MPN^19^ and a low frequency frameshift *CHEK2* mutation has been associated with somatic chromosomal mosaicism^13^. In recent experimental work, suppression of CHEK2 in human cord blood Lin^−^CD34^+^ cells increased cellular proliferation in long term culture. (Bao et al, co-submitted manuscript) These results suggest that while *CHEK2* while may ordinarily limit hematopoietic stem cell expansion, loss of *CHEK2* function may promote self-renewal increasing risk of CHIP.

We next sought to determine whether rare variants in non-coding regions associate with CHIP acquisition. We used a chromatin-state model to predict gene-enhancer pairs in hematopoietic stem cells and aggregated rare variants in these enhancer elements by target gene.^20^ There were 2453 sets of enhancer regions with at least 10 individuals carrying a predicted damaging variant (alpha=2.03 × 10^−5^, see **Methods**). One set of variants in HAPLN1 enhancers exceeded a p-value threshold of p<0.05 after Bonferroni-correction (OR: 6.8, p=1.96 × 10^−5^, **Supplementary Table 10**). *HAPLN1* is an extracellular matrix protein, produced in bone marrow stromal cells that has previously been implicated in NF-κB signaling.^21^

We then asked whether rare non-synonymous coding variants might be associated with clonal fraction when CHIP was present in individuals with single identified driver mutation (N = 4188). There were 19,351 genes where at least 10 individuals had one or more non-synonymous variant (alpha=2.5 × 10^−6^). While no genes exceeded Bonferroni significance, the top gene was Interferon Alpha 1 (*IFNA1*, p= 2.13 × 10^−5^, **Supplementary Table 11-12**).

Lastly, we bioinformatically and experimentally characterized the mechanism by which the non-coding African American-specific variant at the *TET2* locus influenced risk for CHIP. First, iterative conditional analysis at the locus suggested that there was most likely only a single causal variant. Fine-mapping prioritized 25 variants in the credible set (>99% posterior probability), none of which overlaps the coding sequence or promoter of a protein-coding gene. We hypothesized that the causal variant affects an enhancer for *TET2* in hematopoietic stem cells, because heterozygous *Tet2* knockout in mice increases the self-renewal of hematopoietic stem cells *in vivo* and recapitulates the clonal expansion observed in humans with somatic mutations in *TET2*. ^5,9^ Accordingly, we used the Activity-by-Contact (ABC) model to predict which noncoding elements act as enhancers in CD34+ hematopoietic progenitors (see Methods). Only a single variant (rs79901204) in this credible set overlapped an element predicted to regulate any gene, and that element was indeed predicted to regulate TET2 expression. (**Figure 4a, Supplementary Table 13**) To test whether this variant affects enhancer activity, we tested a 600 base pair region containing the regulatory element using a plasmid-based luciferase enhancer assay in CD34+ human hematopoietic progenitor cells (**Figure 4b,c**). The reference sequence activated luciferase expression by 40-fold (versus control constructs with no enhancer sequence), while the T risk allele activated expression by only 10-fold (**Figure 4d**). Indeed, the T risk allele disrupts a consensus GATA/E-Box motif, likely resulting in reduced binding of the activating transcription factor complex GATA1/GATA2. Together, these results suggest that the T risk allele acts to decrease the activity of this enhancer, which in turn reduces expression of *TET2* to promote self-renewal and proliferation of HSPCs. Thus, in this locus both germline noncoding variation and somatic coding variation converge to affect *TET2* and influence the development of CHIP.

**Fig. 4.**
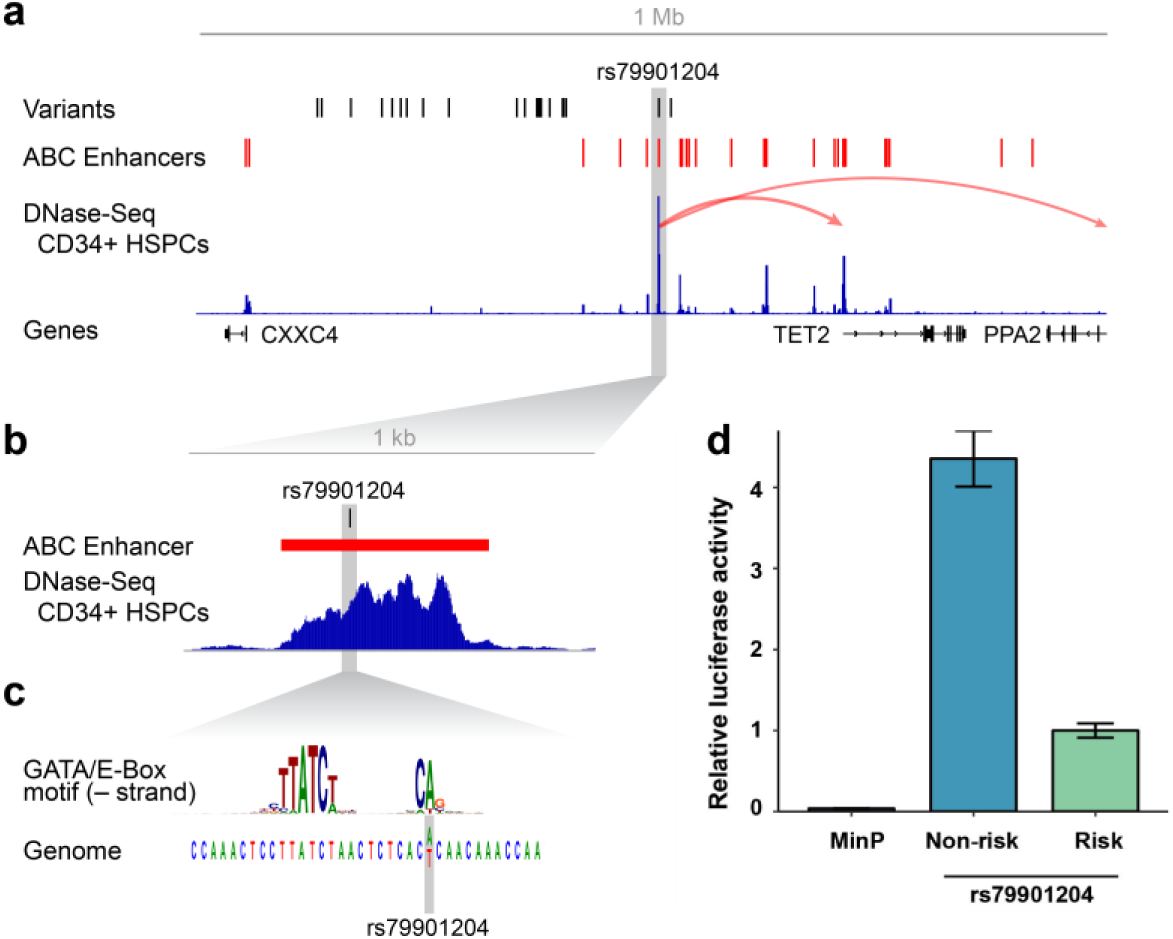
African ancestry specific TET2 locus risk variant disrupts hematopoietic stem cell TET2 enhancer. **a**, the TET2 locus with fine-mapped risk variants, Activity-by-Contact (ABC) hematopoietic stem cell (HSPC) enhancers, DNase-Seq CD34+ HSPC and RefSeq genes. ABC model predicts that rs79901204 disrupts a TET2 enhancer resulting in decreased TET2 expression. **b**, expanded view of TET2 enhancer element. **c**, rs79901204 disrupts a GATA motif/E-Box motif. **d**, luciferase assay in CD34+ primary cells demonstrates four-fold attenuation of enhancer activity by the rs79901204 risk allele relative to the non-risk allele.

Given the role of *TET2* in DNA de-methylation, we hypothesized that carriers of rs79901204 T allele may have differential methylation at the *TET2* locus. We performed a methylation-QTL analysis of a subset of 1592 African Americans and identified significant differential *TET2* locus methylation (**Extended Data Fig. 7, Supplementary Table 14**)

Our observations permit several conclusions. First, our sample size which is nearly an order of magnitude larger than prior CHIP analyses^2,3,12^ enables refinement of CHIP phenotype associations at the level of CHIP driver genes. We find that considerable heterogeneity exists across CHIP phenotypes by driver gene. For example, IL-1b and IL-18 both activate through the inflammasome and increase IL-6.

However, while *TET2* CHIP is associated with increased levels of IL-1b, *JAK2* and *SF3B1* CHIP are associated with IL-18. Second, we identify a convergence of common and rare germline genetic predisposition to leukocyte telomere length, MPN, somatic chromosomal mosaicism and CHIP, suggesting shared causal mechanisms.

Importantly, to date, only CHIP with leukemogenic driver mutations (as opposed to somatic chromosomal mosaicism^13^ or CHIP with unknown driver mutations^12^) has been robustly associated with non-oncologic diseases independently of age. The partially overlapping genetic predisposition we observe across these three clonal phenomena suggest that although there may be similar genetic architecture that predispose individuals to acquiring a somatic mutation, the specific change may be particularly relevant to atherosclerotic disease as opposed to the general phenomenon of clonal hematopoiesis itself.

Important limitations of our study include reduced sensitivity for detecting CHIP with low allele fractions even with high-coverage whole genome sequencing. Ultrasensitive targeted sequencing can facilitate detection of such events but CHIP below the sensitivity of WGS detection (VAF < 5%) may not significantly alter risk for clinical outcomes.^7^ Furthermore, the cross-sectional analyses of CHIP with non-genetic risk factors and biomarkers limit conclusions regarding temporal relationships between CHIP and these features; however, these observations still permit risk prediction for CHIP presence. Notably, inflammatory biomarker analyses are concordant with prior model experiments indicating elevations of observed inflammatory biomarkers as a consequence of CHIP.^5,9^ Lastly, given the age-dependence of CHIP, it is likely that many individuals not observed to have CHIP in this study will develop CHIP in the future.

Overall, comprehensive simultaneous germline and somatic analyses of blood-derived whole genome sequence data demonstrates that germline variation influences the acquisition of somatic mutations in blood cells. Importantly, we anticipate that the TOPMed CHIP dataset defined here will be a valuable tool in establishing associations of CHIP with diverse heart, lung, blood and sleep traits.

## Methods

### Study Samples

Whole genome sequencing (WGS) was performed on 97,691 samples sequenced as part of 52 studies contributing to the NHLBI TOPMed research program as previously described.^7^ Each of the constituent studies provided informed consent on the participating samples. Details on participating cohorts and samples is provided in Supplemental Table S1. The age of participants at time of blood draw was obtained for a subset of 82,807 of the samples. The median age was 55, the mean age 52.5, and the maximum age 98. The age distribution varied across the constituent cohorts (Supplemental Table S1).

### WGS Processing, Variant Calling and CHIP annotation

BAM files were remapped and harmonized through a previously described unified protocol.^23^ SNPs and short indels were jointly discovered and genotyped across the TOPMed samples using the GotCloud pipeline.^24^ An SVM filter was trained to discriminate between true variants and low-quality sites. Sample quality was assessed through pedigree errors, contamination estimates, and concordance between self-reported sex and genotype inferred sex. Variants were annotated using snpEff 4.3.

Putative somatic SNPs and short indels were called with GATK Mutect2 (https://software.broadinstitute.org/gatk). Briefly, Mutect2 searches for sites where there is evidence for variation, and then performs local reassembly. It uses an external reference of recurrent sequencing artifacts termed a “panel of normal” to filter out these sites, and calls variants at sites where there is evidence for somatic variation. An external reference of germline variants^25^ was provided to filter out likely germline calls. We deployed this variant calling process on Google Cloud using Cromwell (https://github.com/broadinstitute/cromwell). The caller was run individually for each sample with the same settings. The Cromwell WDL configuration file is available from the authors upon request.

Samples were annotated as having CHIP if the Mutect2 output contained one or more of a pre-specified list of putative CHIP variants as previously described^2,5^ (Supplemental Table S2).

### Blood traits

Conventionally measured blood cell counts and indices were selected for analysis including: hemoglobin, hematocrit, red blood cell count, white blood cell count, basophil count, eosinophil count, neutrophil count, lymphocyte count, monocyte count, platelet count, mean corpuscular hemoglobin, mean corpuscular hemoglobin concentration, mean corpuscular volume, mean platelet volume and red cell distribution width. Phenotypes were collected by each cohort, centrally harmonized by the TOPMed Data Coordinating Center (DCC). Additional documentation about harmonization algorithms for each specific trait is available from the TOPMed DCC and accompanies the data on the dbGaP TOPMed Exchange area. Up to 37,653 individuals from 10 cohorts where utilized for this analysis that had one or more blood traits measured concurrently or following the blood draw used for CHIP ascertainment. Traits were first log2 normalized and then analyzed using a general linear regression model with CHIP status, age, sex, study and the first 10 ancestry principal components as covariates.

### Lipid phenotypes

Conventionally measured plasma lipids, including total cholesterol, LDL-C, HDL-C, and triglycerides, were included for analysis. LDL-C was either calculated by the Friedewald equation when triglycerides were <400 mg/dl or directly measured. Given the average effect of statins, when statins were present, total cholesterol was adjusted by dividing by 0.8 and LDL-C by dividing by 0.7. Triglycerides were natural log transformed for analysis. Phenotypes were harmonized by each cohort and deposited into dbGaP TOPMed Exchange area as previously described.^26^ Up to 28,310 individuals from 19 cohorts where utilized for this analysis that had one or more lipid trait measured concurrently or following the blood draw used for CHIP ascertainment. Lipid traits were first normalized for age, sex and ancestry principal components and then analyzed using a general linear regression model with CHIP status, age, sex, study and the first 10 ancestry principal components as covariates.

### Inflammatory Markers

A set of makers previously implicated in mediating cardiometabolic disease were analyzed including: CD-40, CRP, E-Selectin, ICAM-1, IL-1b, IL-6, IL-10, IL-18, 8-epi PGF2a, Lp-PLA2 mass and activity, MCP1, MMP9, MPO, OPG, P-selectin, TNF-Alpha, TNF-Alpha Receptor 1, TNF-receptor 2. Phenotypes were collected by each cohort, centrally harmonized by the TOPMed DCC and then deposited into dbGaP TOPMed Exchange area. Additional documentation about harmonization algorithms for each specific trait is available from the TOPMed DCC and accompanies the data on dbGaP. Up to 22,092 individuals from 10 cohorts where utilized for this analysis that had one or more inflammatory marker measured concurrently or following the blood draw used for CHIP ascertainment. Inflammatory markers were first normalized using a log2(x+1) transformation and then analyzed using a general linear regression model with CHIP status, age, sex, study and the first 10 ancestry principal components as covariates.

### Single Variant Association

Single variant association for each variant in Freeze 8 with MAF > 0.1% and MAC > 20 was performed with SAIGE^23^, and analysis was performed using the TOPMed Encore analysis server (https://encore.sph.umich.edu). CHIP driver status was dichotomized into a case-control phenotype based on the presence of at least one driver mutation. Prior to running single variant association tests, a logistic mixed model was fit using the lme4 R package^24^ to estimate the probability of the CHIP case control status conditional on a spline transformation of the centered age, genotype inferred sex, and cohort. The cohort was included as a random intercept which represents study specific contributions to the log-odds of CHIP at the mean sample age. Age was modeled with a spline to capture the non-linearity of the relationship between age and CHIP. This model was chosen over comparable models based on its AIC. Combining the age, inferred sex, and study into a single quantity aided the convergence of SAIGE compared to the inclusion of these terms separately. The first 10 principal components were also included as covariates.

Given that CHIP is unlikely to manifest in younger individuals, these individuals are effectively censored in our analysis set – that is, a young individual that does not presently have CHIP may still develop CHIP in the future. To avoid the power loss associated with misclassification of controls, we pruned these individuals from our analysis set. The single variant association analysis was run on a pruned set of samples that excluded those which had less than a 1% probability CHIP as estimated by the aforementioned model. This excluded 21,712 samples leading to a final analysis set of 65,405 which was used for downstream association analyses.

### Fine mapping

We applied FINEMAP to the summary statistics from SAIGE, using the z-score and LD matrices as input. We fine-mapped the TET2 locus using the summary statistics from the African ancestry single variant summary statistics and estimated LD on the same set of samples using plink. We set the maximum number of causal SNPs in the region to 10 and used a shotgun stochastic search.

### Rare Variant Analyses

Collapsing burden tests were applied to specific variant grouping schemes using EPACTS. The same covariates as the single variant tests were used on the same set of samples. We used burden tests due to their limited compute requirements, which were considerable for the number of variants and samples tested. Two grouping schemes were specified: the first groups coding variation, and the second groups putative regulatory elements in a relevant cell line. The first used all putative LOF variants as identified by snpEff. Given that some variants were present in both the Mutect2 calls and the germline variant calls, we pruned the LOF variants to exclude variants that were present in both call sets. The second grouping scheme used all variants in regions that were predicted enhancers for CD34 cells that had CADD scores of at least 10. Predicted enhancers were identified by the activity-by-contact model.^20^

### Predicting enhancer-gene regulation for TET2

We used the Activity-by-Contact (ABC) model to predict which enhancers regulate which genes in CD34+ hematopoietic progenitor cells as previously described^20^, with minor modifications as follows.

Briefly, this model predicts the effect of each putative regulatory element (defined as a DNase peak within 5Mb of a given promoter) by multiplying the Activity of each element (estimated from DNase-seq and H3K27ac ChIP-seq) by its Contact with a target promoter (estimated from Hi-C data). The ABC score of a single element on a gene’s expression is the predicted effect of that element divided by the sum of the predicted effects of all elements for a given gene.

We identified putative regulatory elements by using MACS2 to call peaks in DNase-seq data from mobilized CD34+ hematopoietic progenitor cells from the Roadmap Epigenome Project (downloaded from http://egg2.wustl.edu/roadmap/data/byFileType/alignments/consolidated/E050-DNase.tagAlign.gz) Initially we considered all peaks with p-value < 0.1. To further refine this list, we kept the 100,000 peaks with the highest number of DNase-seq reads. We then resized these peaks to be 500 bp in length centered on the peak summit, merging any overlapping peaks, and removed any peaks overlapping ENCODE “blacklisted regions”^27^ (regions of the genome previously observed to accumulate anomalous numbers of reads in epigenetic sequencing experiments; downloaded from https://sites.google.com/site/anshulkundaje/projects/blacklists). To this peak list, we added 500 bp regions centered on the transcription start site of all genes. Any overlapping regions resulting from these additions or extensions were merged.

Within each putative regulatory element, we estimated enhancer Activity as the geometric mean of read counts from DNase-seq and H3K27ac ChIP-seq data from the Roadmap Epigenome Project (http://egg2.wustl.edu/roadmap/data/byFileType/alignments/consolidated/E050-DNase.tagAlign.gz, and E050-H3K27ac.tagAlign.gz).

We estimated enhancer-promoter Contact from the KR-normalized Hi-C contact maps in primary CD34+ cells. we then calculated effect of each putative enhancer-gene connection by multiplying the Activity and Contact for that element and gene. Dividing the effect of each element by the sum of effects for all elements for a given gene yields the ABC score:

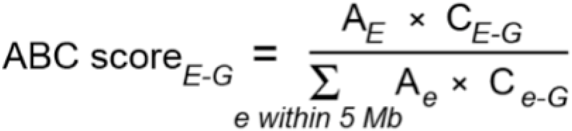

To call predicted enhancer-gene connections, we used a threshold on the ABC score of 0.015. The rs79901204 variant overlapped an enhancer with ABC score of 0.0308 for TET2, which, based on comparison of ABC scores to large-scale enhancer perturbation datasets, corresponds to a positive predictive value of approximately 61%.

### Functional Evaluation of TET2 locus

The genomic region containing risk and non-risk allele of the variant rs79901204 (600bp) was synthesized as gblocks (IDT Technologies) and cloned into the Firefly luciferase reporter constructs (pGL4.24) using NheI and EcoRV sites. The Firefly constructs (500ng) were co-transfected with pRL-SV40 Renilla luciferase constructs (50ng) into 100,000 K562 cells using Lipofectamine LTX (Invitrogen) according to manufacturer’s protocols. Cells were harvested after 48 hours and the luciferase activity measured by Dual-Glo Luciferase Assay system (Promega).

### Methylation-QTL analysis of TET2 locus

Illumina MethylationEPIC 850K array data interrogating over 850,000 CpG DNA methylation sites was generated at the University of Washington’s Northwest Genomic Center from blood samples collected from African Americans at the Jackson Heart Study baseline exam. Fluorescent signal intensities were preprocessed with the R package *minfi*^28^ using the normal-exponential out-of-band (noob) background correction method with dye-bias normalization. N = 1756 total samples (1203 women and 653 men) remained after severe outliers were identified and removed. Methylation levels at each CpG site were then quantified as β values, defined as the ratio of intensities between methylated (M) and unmethylated (U) signals where β = M/(M+U+100). Values therefore ranged from β = 0 (completely unmethylated) to β = 1 (completely methylated). Batch correction for assay plate position was performed on the β values via *ComBat.*^29^ Relative leukocyte cell counts (CD8+ T-lymphocytes, CD4+ T-lymphocytes, Natural Killer cells, B cells, Monocytes, and Granulocytes) were estimated as previously described by Houseman^30^ and Horvath^31^.

To investigate local methylation in the CXXC4/TET2 locus, a region of interest +/-1Mb of the variant rs79901204 was considered containing a total of 311 CpG sites for analysis. The analysis included a subset of 1587 African Americans from the Jackson Heart Study. Of these individuals, 48 had CHIP and 103 were carriers of the rs79901204 variant. A linear mixed effects model was fitted using *CpGassoc*^32^ in R 3.6.0 with rs79901204 as the predictor and the batch-corrected methylation β levels as the dependent variable, adjusting for age, sex, estimated cell counts, and CHIP status. A Bonferroni corrected threshold of *P* = 1.6 × 10^−4^ was used to establish statistical significance.

## Supporting information

Supplemental Acknowledgements

## DATA AVAILABILITY

Individual whole-genome sequence data for TOPMed whole genomes, individual-level harmonized phenotypes and the CHIP variant call sets used in this analysis are available through restricted access via the dbGaP TOPMed Exchange Area available to TOPMed investigators. Controlled-access release to the general scientific community via dbGaP is ongoing. Accession numbers for these datasets are: phs001237, phs000951, phs001416, phs001515, phs000974, phs001644, phs001211, phs000964, phs001612, phs001467, phs000988, phs001368, phs000920, phs001468, phs001387, phs001446, phs001217, phs001215, phs001293, phs001218, phs001395, phs001472, phs000921, phs001402, phs000972, phs001624, phs001345, phs001032, phs000956, phs001062, phs001726, phs001143, phs000954, phs001359, phs001466, phs001207, phs000993, phs001661, phs001607, phs001542, phs001608, phs001545, phs001601, phs001412, phs001189, phs001598, phs001543, phs001725, phs000946, phs001435, phs000997, phs001434, phs001546. Summary-level genotype data are available through the BRAVO browser (https://bravo.sph.umich.edu/).

## ACKNOWLEDGEMENTS

Investigators who conducted this research report individual research support from R35 HL135818 (S. Redline), P01 HL132825 (S. Weiss), R01HL091357 and R01HL055673 (D. Arnett), W81XWH-17-1-0597 (D. Schwartz), K01 HL135405 (B. Cade), P01 HL132825 (J Lasky-Su), K01HL136700 (S. Aslibekyan), R01HL113323 (J Curran), R01HL1333040 (D. Weeks), 1R01HL138737 (D. Darbar), P01 HL132825 (P. Kachroo), T32 HL129982 (L. Raffield), R01HL113323 (J. Blangero), HHS-N268201800002I (T. Blackwell and A Smith), U54GM115428 (J. Wilson), Claudia Adams Barr Program for Innovative Cancer Research (V. Sankaran), R01142711, MGH Hassenfeld Scholar Award (P. Natarajan), Fondation Leducq TNE-18CVD04 (B. Ebert, P. Natarajan, S. Kathiresan), 5UM1HG008895 and the Ofer and Shelly Nemirovsky Research Scholar Award from Massachusetts General Hospital (S. Kathiresan).

Whole genome sequencing (WGS) for the Trans-Omics in Precision Medicine (TOPMed) program was supported by the National Heart, Lung and Blood Institute (NHLBI). Centralized read mapping and genotype calling, along with variant quality metrics and filtering were provided by the TOPMed Informatics Research Center (3R01HL-117626-02S1; contract HHSN268201800002I). Phenotype harmonization, data management, sample-identity QC, and general study coordination were provided by the TOPMed Data Coordinating Center (3R01HL-120393-02S1; contract HHSN268201800001I). We gratefully acknowledge the studies and participants who provided biological samples and data for TOPMed.

---

**Extended Data Fig. 1.**
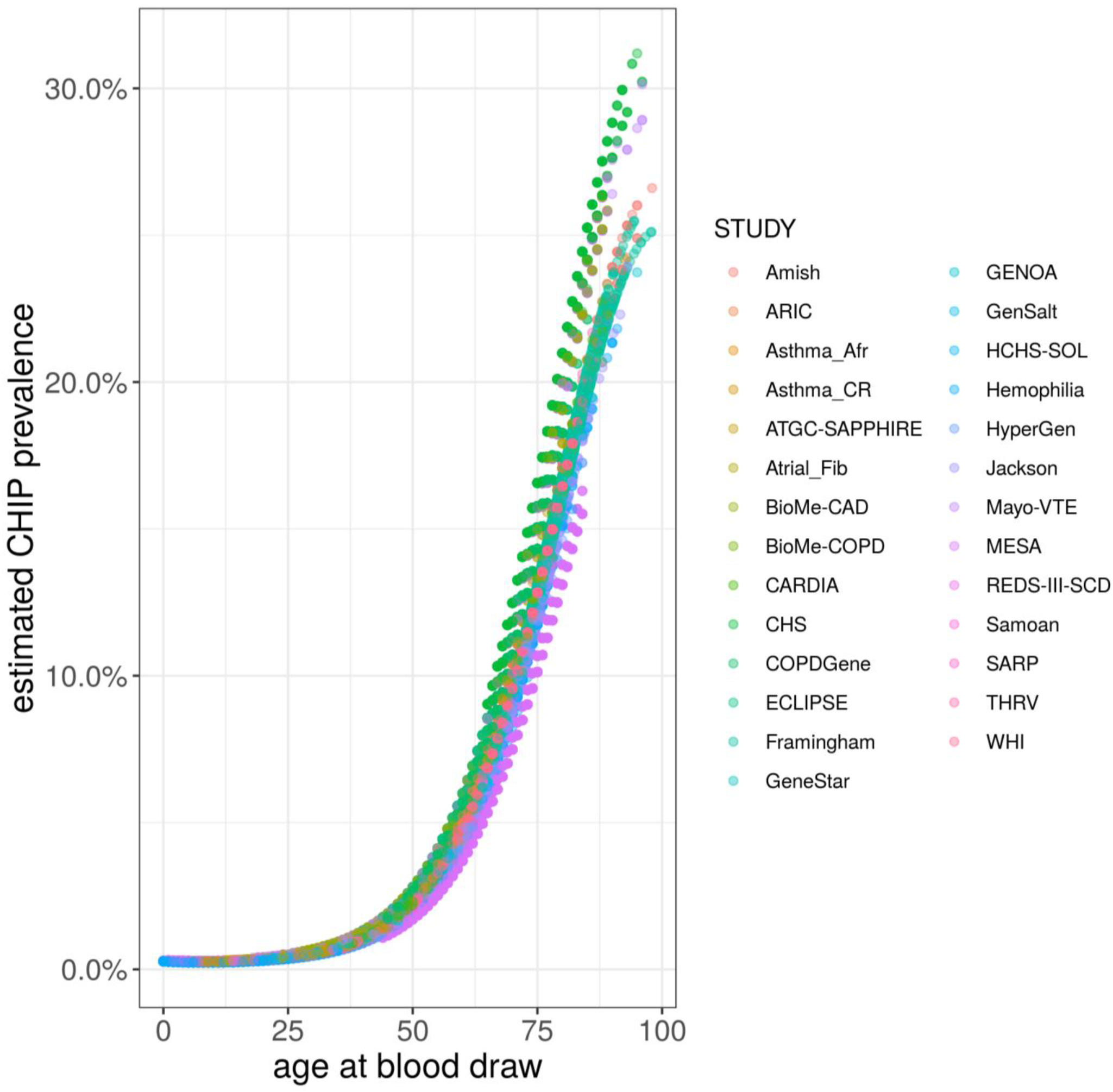
CHIP prevalence by study, CHIP prevalence with age at blood draw was highly concordant across sequenced cohorts

**Extended Data Fig. 2.**
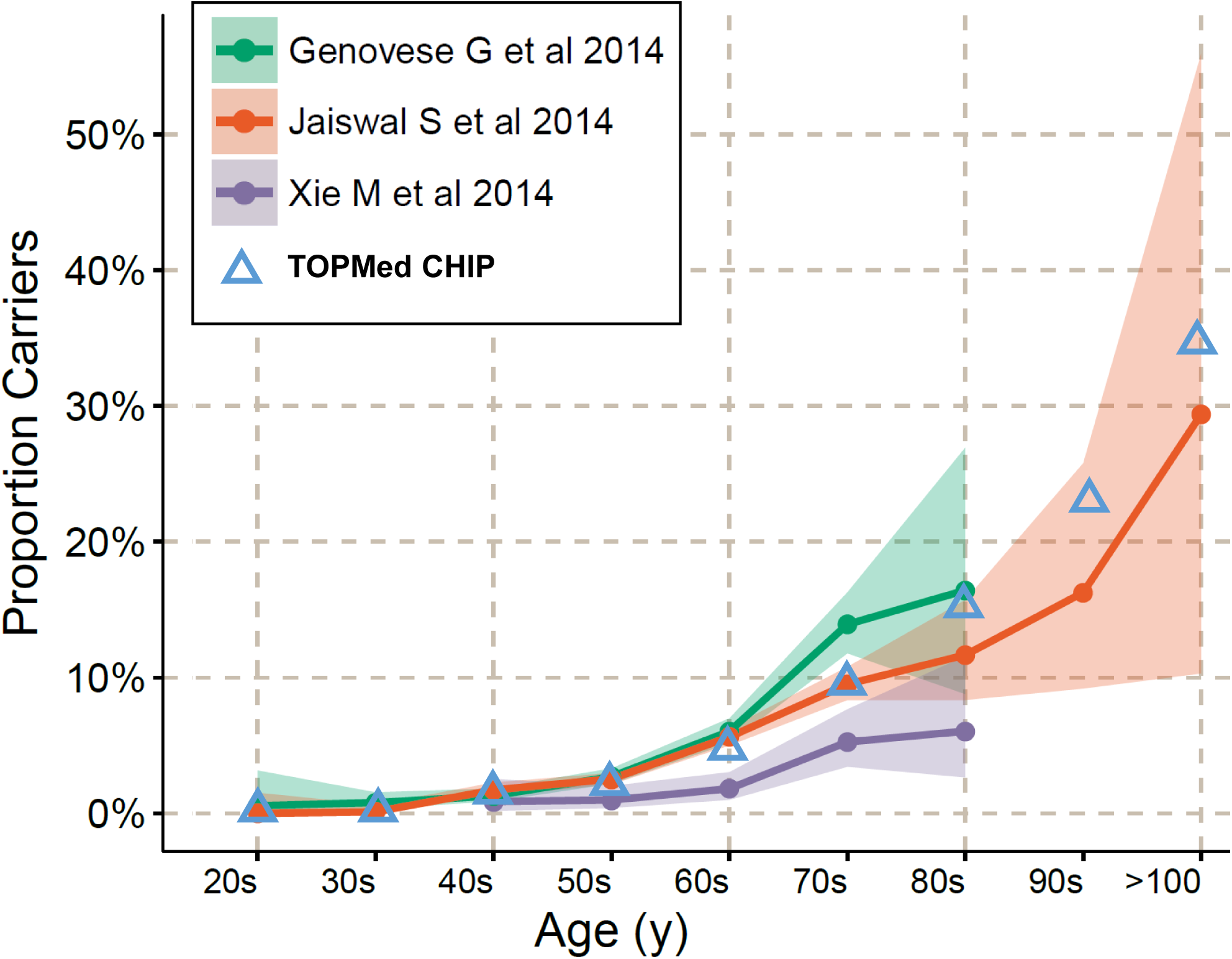
CHIP prevalence in comparison to prior reports, CHIP prevalence with age in this study (blue triangles) was highly consistent with previously observed CHIP prevalence.

**Extended Data Fig. 3.**
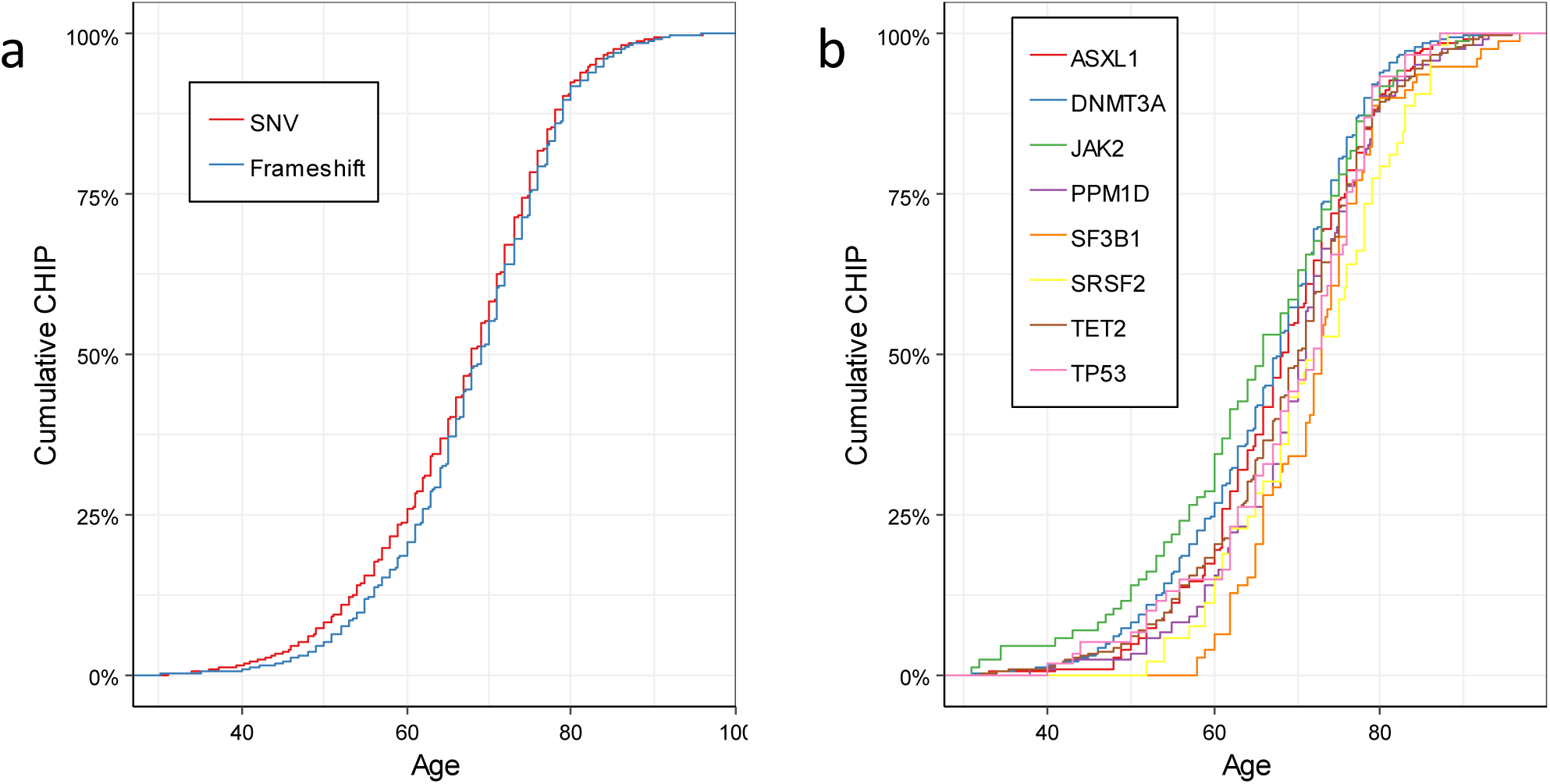
CHIP age association by mutational mechanism and gene. **a**, cumulative density plot of CHIP incidence with age stratified by single nucleotide variant (SNV) vs frameshift mutations. SNVs were observed in younger individuals than Frameshift mutations (Wilcox rank sum test: p=0.01). **b**, cumulative density plot of CHIP incidence with age stratified by driver gene

**Extended Data Fig. 4.**
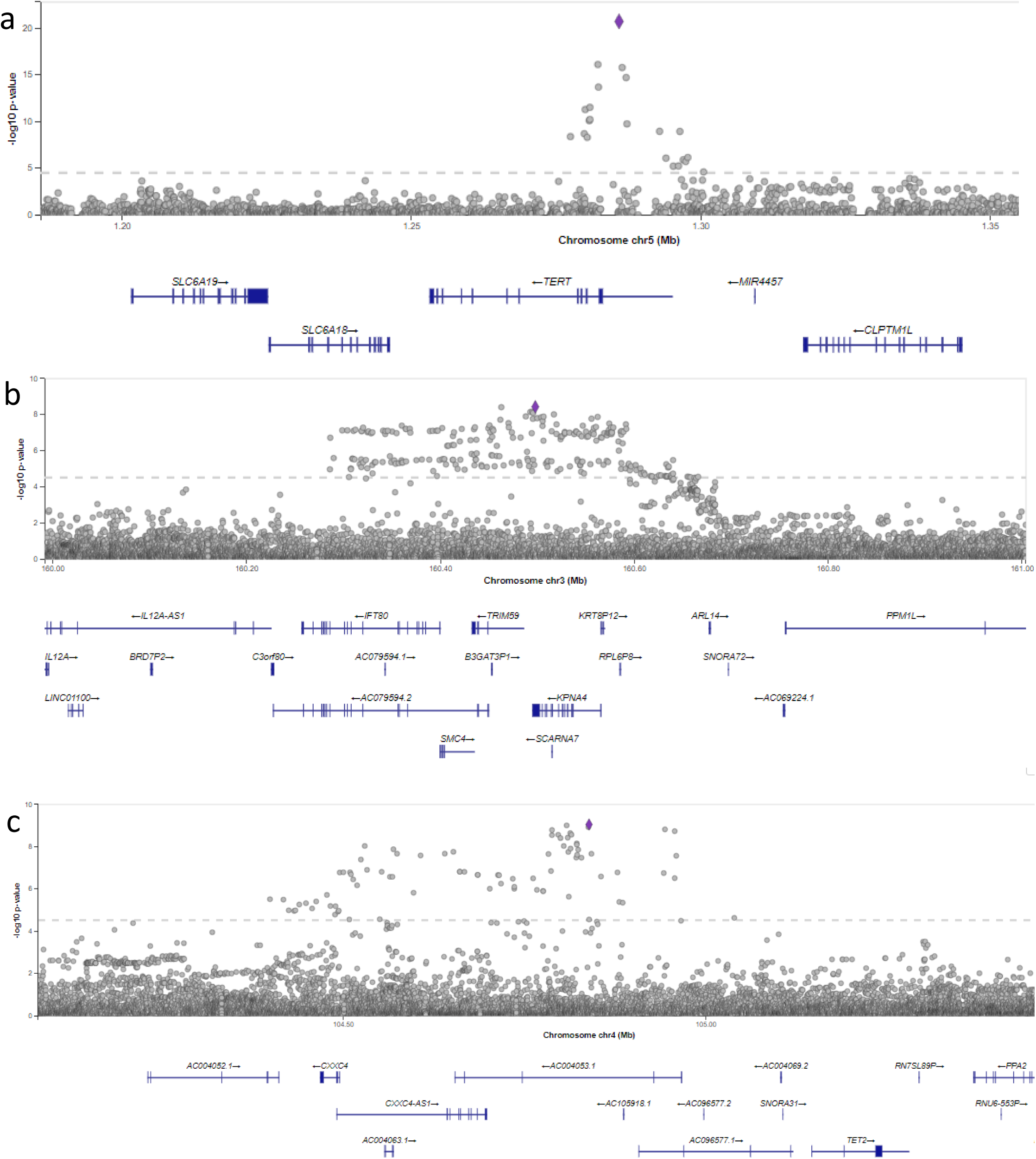
CHIP Single variant association regional association plots. **a**, TERT locus **b**, TRIM59/KPNA4 locus **c**, TET2 locus

**Extended Data Fig. 5.**
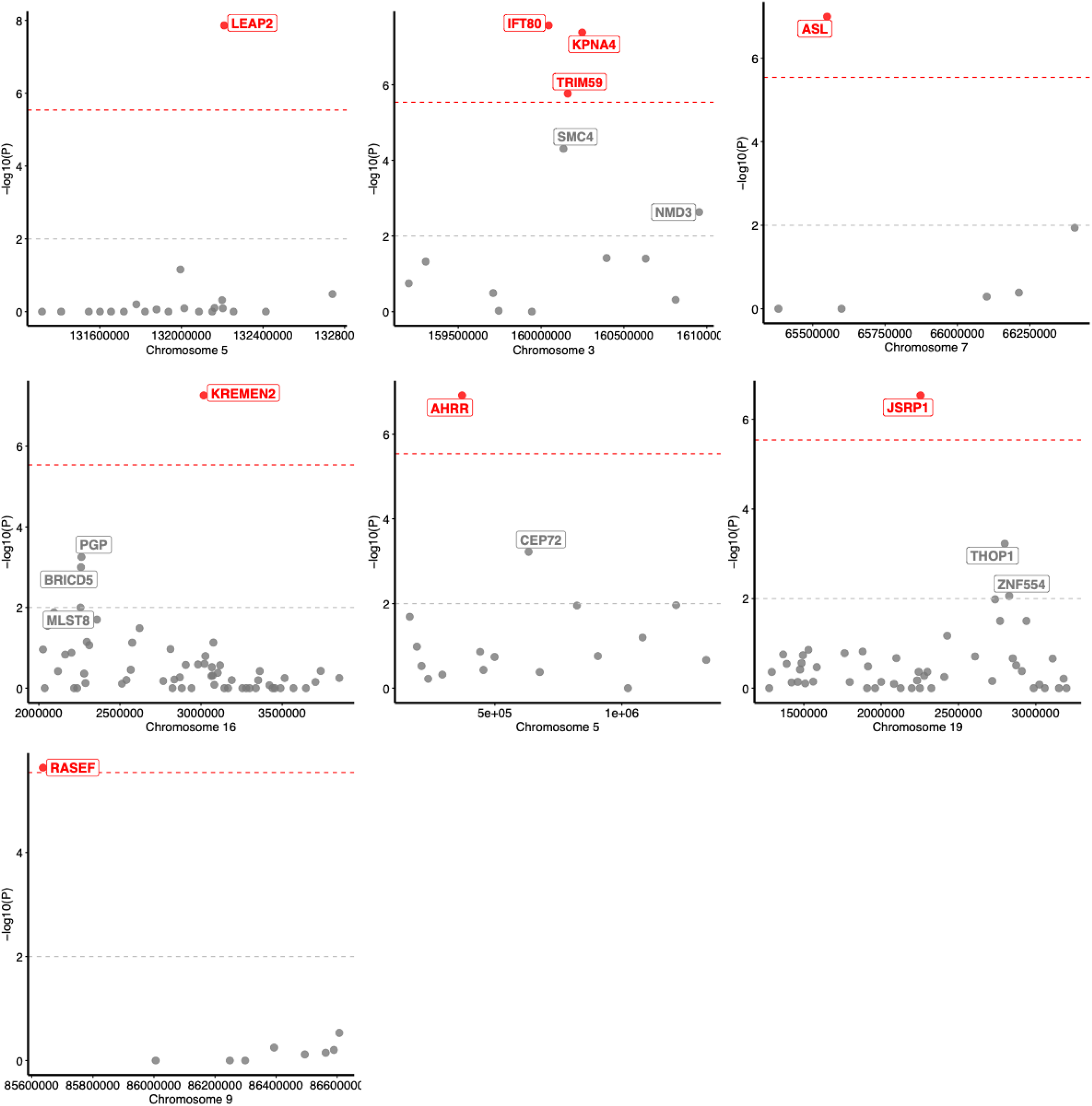
UTMOST combined CHIP TWAS results across 48 tissues identified 7 significant loci (p<2.9 × 10^−6^)

**Extended Data Fig. 6.**
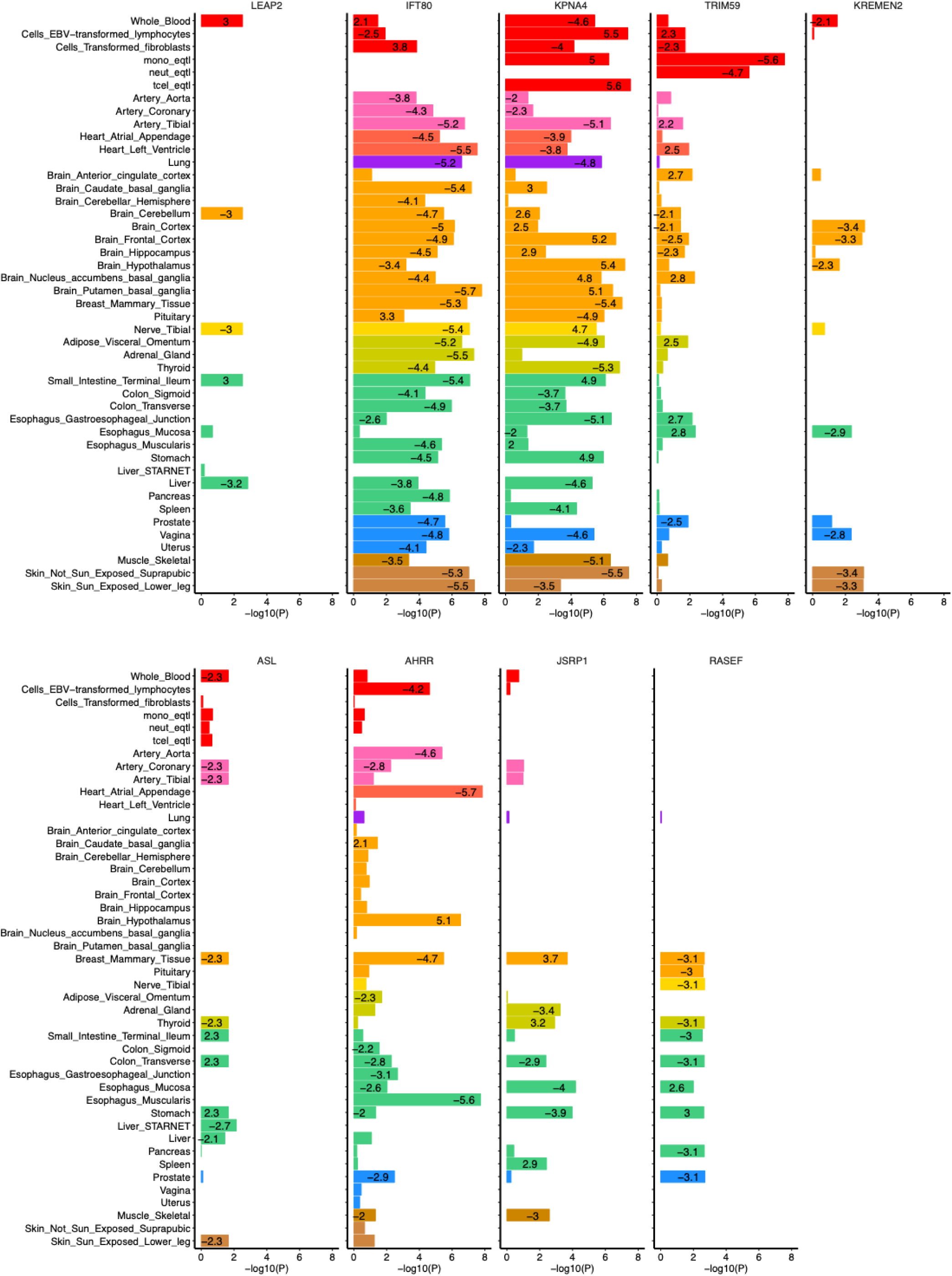
Tissue-specific results from the top 9 overall UTMOST-significant genes. eQTL z-scores for associations with P<0.05 are displayed in each bar

**Extended Data Fig. 7.**
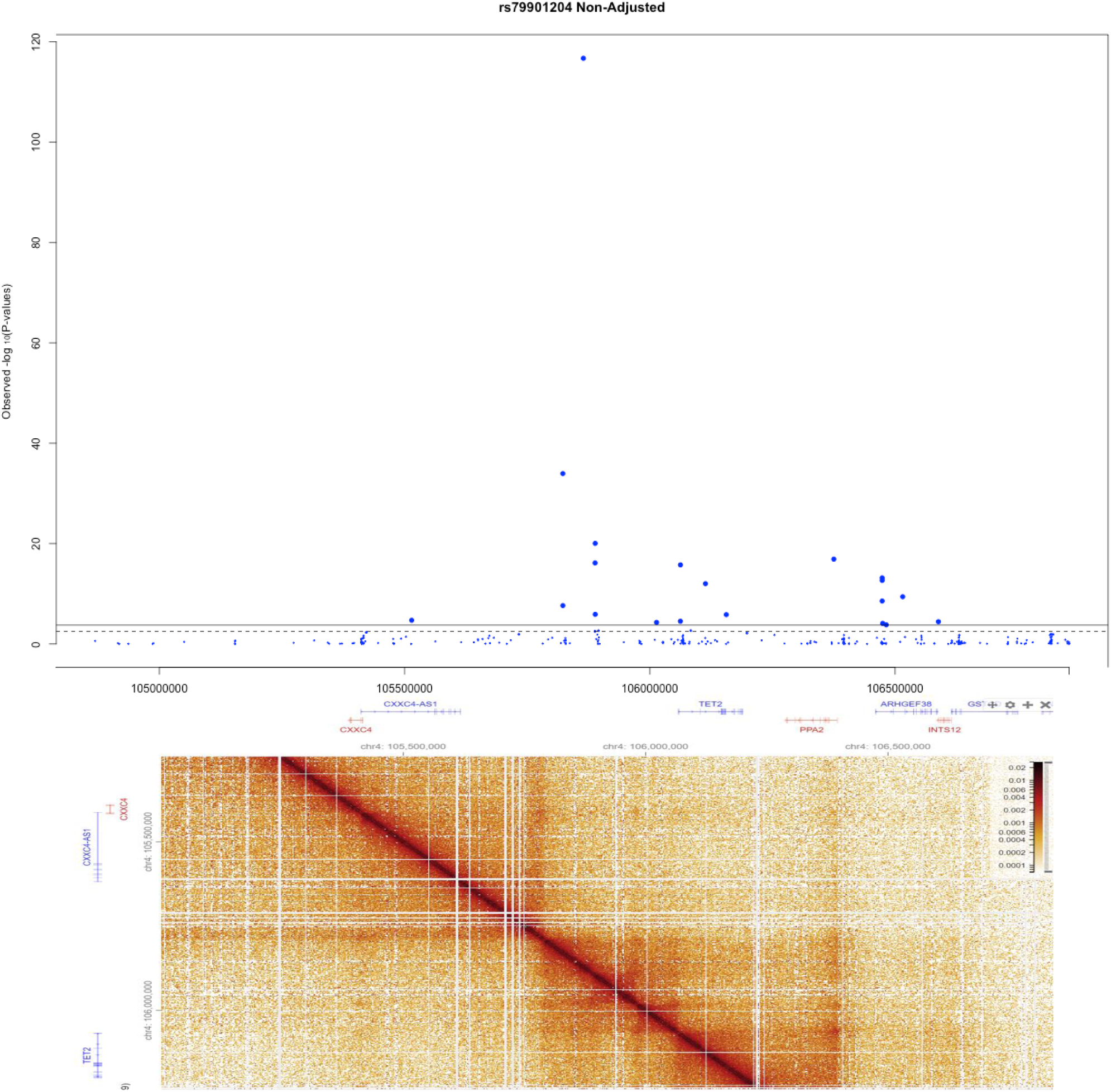
African ancestry specific TET2 locus risk variant rs79901204 associated with altered TET2 locus methylation. **top**, Methylation Quantitative Trait association of rs79901204 variant with cpg methylation probes in the TET2 locus demonstrate that carriers of rs79901204 have an altered peripheral leukocyte methylation profile most notably for the *TET2* gene as well as for near by genes *ARHGEF38* and *PPA2*. **bottom**, Hi-C domains from GM12878 (GEO ID: GSM1551688) visualized with higlass.io

