## Supplemental Acknowledgements for "Inherited Causes of Clonal Hematopoiesis of Indeterminate Potential in TOPMed Whole Genomes"

**NHLBI TOPMed: Sequencing Acknowledgements**

| **TOPMed Accession #** | **Parent Study Short Name** | **Parent Study Full Name** | **TOPMed Phase** | **TOPMed Project** | **Omics Center** | **Omics Support Grant/Contract Number** |
| --- | --- | --- | --- | --- | --- | --- |
| phs001543 | AFLMU | Atrial Fibrillation Biobank Ludwig Maximilian University Study | 2.5 | AFGen | BROAD | HHSN268201500014C |
| phs000956 | Amish | Genetics of Cardiometabolic Health in the Amish | CCDG co-funded | AFGen | BROAD | 3R01HL121007-01S1 |
| phs001543 | AFLMU | Atrial Fibrillation Biobank Ludwig Maximilian University Study | 1 | Amish | BROAD | 3UM1HG008895-01S2 |
| phs001211 | ARIC | Atherosclerosis Risk in Communities Study | 1 | AFGen | BROAD | 3R01HL092577-06S1 |
| phs001211 | ARIC | Atherosclerosis Risk in Communities Study VTE cohort | 2 | VTE | BAYLOR | 3U54HG003273-12S2, HHSN268201500015C |
| phs001435 | AustralianFamilialAF | Molecular Mechanisms of Inherited Cardiomyopathies and Arrhythmias in the Australian Familial AF Study | 1.5 | AFGen | BROAD | 3U54HG003067-12S2, 3U54HG003067-13S1 |
| phs001143 | BAGS | New Approaches for Empowering Studies of Asthma in Populations of African Descent - Barbados Asthma Genetics Study | 1 | BAGS | ILLUMINA | 3R01HL104608-04S1 |
| phs001644 | BioMe | Mount Sinai BioMe Biobank | 3 | BioMe | BAYLOR | HHSN268201600033I |
| phs001644 | BioMe | Mount Sinai BioMe Biobank | 3 | BioMe | WASHU MGI | HHSN268201600037I |
| phs001644 | BioMe | Mount Sinai BioMe Biobank | CCDG co-funded | AFGen | WASHU MGI | 3UM1HG008853-01S2 |
| phs001624 | BioVU_AF | Mount Sinai BioMe Biobank | CCDG co-funded | AFGen | BAYLOR | 3UM1HG008898-01S3 |
| phs001726 | CAMP | Childhood Asthma Management Program | 3 | CRA_CAMP | UW NWGC | HHSN268201600032I |
| phs001612 | CARDIA | Coronary Artery Risk Development in Young Adults | 3 | CARDIA | BAYLOR | HHSN268201600033I |
| phs001189 | CCAF | Cleveland Clinic Atrial Fibrillation Study | 1 | AFGen | BROAD | 3R01HL092577-06S1 |
| phs000954 | CFS | Cleveland Family Study - WGS Collaboration | 1 | CFS | UW NWGC | 3R01HL098433-05S1 |
| phs000954 | CFS | Cleveland Family Study - WGS Collaboration | 3.5 | CFS | UW NWGC | HHSN268201600032I |
| phs001368 | CHS | Cardiovascular Health Study | 2 | VTE | BAYLOR | 3U54HG003273-12S2, HHSN268201500015C |
| phs001368 | CHS | Cardiovascular Health Study | 3 | CHS | BAYLOR | HHSN268201600033I |
| phs000951 | COPDGene | Genetic Epidemiology of COPD Study | 1 | COPD | UW NWGC | 3R01HL089856-08S1 |
| phs000951 | COPDGene | Genetic Epidemiology of COPD Study | 2 | COPD | BROAD | HHSN268201500014C |
| phs000951 | COPDGene | Genetic Epidemiology of COPD Study | 2.5 | COPD | BROAD | HHSN268201500014C |
| phs000988 | CRA | The Genetic Epidemiology of Asthma in Costa Rica - Asthma in Costa Rica cohort | 1 | CRA_CAMP | UW NWGC | 3R37HL066289-13S1 |
| phs000988 | CRA | The Genetic Epidemiology of Asthma in Costa Rica - Asthma in Costa Rica cohort | 3 | CRA_CAMP | UW NWGC | HHSN268201600032I |
| phs001546 | DECAF | Determining the association of chromosomal variants with non-PV triggers and ablation-outcome in DECAF | 1.5 | AFGen | BROAD | 3U54HG003067-12S2, 3U54HG003067-13S1 |
| phs001472 | ECLIPSE | Evaluation of COPD Longitudinally to Identify Predictive Surrogate End-points | 3 | COPD | WASHU MGI | HHSN268201600037I |
| phs000946 | EOCOPD | Boston Early-Onset COPD Study | 1 | COPD | UW NWGC | 3R01HL089856-08S1 |
| phs000974 | FHS | Framingham Heart Study | 1 | AFGen | BROAD | 3R01HL092577-06S1 |
| phs000974 | FHS | Framingham Heart Study | 1 | FHS | BROAD | 3U54HG003067-12S2 |
| phs001542 | GALAI | ATGC Gene-Environment, Admixture and Latino Asthmatics Study I Asthma | 3 | ATGC | UW NWGC | HHSN268201600032I |
| phs000920 | GALAII | Gene-Environment, Admixture and Latino Asthmatics Study | 1 | PGX_Asthma | NYGC | 3R01HL117004-02S3 |
| phs000920 | GALAII | ATGC Gene-Environment, Admixture and Latino Asthmatics Study II Asthma | 3 | ATGC | UW NWGC | HHSN268201600032I |
| phs001218 | GeneSTAR | Genetic Studies of Atherosclerosis Risk | 2 | AA_CAC | BROAD | HHSN268201500014C |
| phs001218 | GeneSTAR | Genetic Studies of Atherosclerosis Risk | 2 | GeneSTAR | MACROGEN | 3R01HL112064-04S1 |
| phs001218 | GeneSTAR | Genetic Studies of Atherosclerosis Risk | legacy | GeneSTAR | ILLUMINA | R01HL112064 |
| phs001345 | GENOA | Genetic Epidemiology Network of Arteriopathy | 2 | AA_CAC | BROAD | HHSN268201500014C |
| phs001345 | GENOA | Genetic Epidemiology Network of Arteriopathy | 2 | HyperGEN_GENOA | UW NWGC | 3R01HL055673-18S1 |
| phs001217 | GenSalt | Genetic Epidemiology Network of Salt Sensitivity | 2 | GenSalt | BAYLOR | HHSN268201500015C |
| phs001725 | GGAF | Groningen Genetics of Atrial Fibrillation Study | CCDG co-funded | AFGen | BAYLOR | 3UM1HG008898-01S3 |
| phs001359 | GOLDN | Genetics of Lipid Lowering Drugs and Diet Network | 2 | GOLDN | UW NWGC | 3R01HL104135-04S1 |
| phs001395 | HCHS_SOL | Hispanic Community Health Study - Study of Latinos | 3 | HCHS_SOL | BAYLOR | HHSN268201600033I |
| phs000993 | HVH | Heart and Vascular Health Study | 1 | AFGen | BROAD | 3R01HL092577-06S1 |
| phs000993 | HVH | Heart and Vascular Health Study | 2 | VTE | BAYLOR | 3U54HG003273-12S2, HHSN268201500015C |
| phs001293 | HyperGEN | Hypertension Genetic Epidemiology Network | 2 | HyperGEN_GENOA | UW NWGC | 3R01HL055673-18S1 |
| phs001545 | INSPIRE_AF | Intermountain Heart Study | CCDG co-funded | AFGen | BROAD | 3UM1HG008895-01S2 |
| phs001607 | IPF | Whole Genome Sequencing in Familial and Sporadic Idiopathic Pulmonary Fibrosis | 3 | IPF | WASHU MGI | HHSN268201600037I |
| phs000964 | JHS | Jackson Heart Study | 1 | JHS | UW NWGC | HHSN268201100037C |
| phs001598 | JHU_AF | The Johns Hopkins University School of Medicine Atrial Fibrillation Genetics Study | CCDG co-funded | AFGen | BROAD | 3UM1HG008895-01S2 |
| phs001402 | Mayo_VTE | Mayo Clinic Venous Thromboembolism Study | 2 | VTE | BAYLOR | 3U54HG003273-12S2, HHSN268201500015C |
| phs001416 | MESAFam | Multi-Ethnic Study of AtherosclerosisMulti-Ethnic Study of Atherosclerosis Family cohort | 2 | AA_CAC | BROAD | HHSN268201500014C |
| phs001416 | MESA | Multi-Ethnic Study of Atherosclerosis | 2 | MESA | BROAD | 3U54HG003067-13S1 |
| phs001062 | MGH_AF | Massachusetts General Hospital Atrial Fibrillation Study | 1 | AFGen | BROAD | 3R01HL092577-06S1 |
| phs001062 | MGH_AF | Massachusetts General Hospital Atrial Fibrillation Study | 1.4 | AFGen | BROAD | 3U54HG003067-12S2, 3U54HG003067-13S1 |
| phs001062 | MGH_AF | Massachusetts General Hospital Atrial Fibrillation Study | 1.5 | AFGen | BROAD | 3U54HG003067-12S2, 3U54HG003067-13S1 |
| phs001062 | MGH_AF | Massachusetts General Hospital Atrial Fibrillation Study | CCDG co-funded | AFGen | BROAD | 3UM1HG008895-01S2 |
| phs001434 | miRhythm | Defining time-dependent genetic and transcriptomic responses to cardiac injury among patients with arrhythmias | 1.5 | AFGen | BROAD | 3U54HG003067-12S2, 3U54HG003067-13S1 |
| phs001515 | MLOF | My Life, Our Future: Genotyping for Progress in Hemophilia | 2 | MLOF | NYGC | HHSN268201500016C |
| phs001515 | MLOF | My Life, Our Future: Genotyping for Progress in Hemophilia | 3 | MLOF | BAYLOR | HHSN268201600033I |
| phs001608 | OMG_SCD | Outcome Modifying Genes in Sickle Cell Disease | 2 | OMG_SCD | BAYLOR | HHSN268201500015C |
| phs001466 | PharmHU | The Pharmacogenomics of Hydroxyurea in Sickle Cell Disease | 2 | PharmHU | BAYLOR | HHSN268201500015C |
| phs001601 | PMBB_AF | Early-onset Atrial Fibrillation in the Penn Medicine BioBank Cohort | CCDG co-funded | AFGen | WASHU MGI | 3UM1HG008853-01S2 |
| phs001601 | PMBB_AF | Early-onset Atrial Fibrillation in the Penn Medicine BioBank Cohort | CCDG co-funded | AFGen | BROAD | 3UM1HG008895-01S2 |
| phs001468 | REDS-III_Brazil | Recipient Epidemiology and Donor Evaluation Study-III | 2 | REDS-III_Brazil | BAYLOR | HHSN268201500015C |
| phs001215 | SAFS | Whole Genome Sequencing to Identify Causal Genetic Variants Influencing CVD Risk - San Antonio Family Studies | 1 | SAFS | ILLUMINA | 3R01HL113323-03S1 |
| phs001215 | SAFS | Whole Genome Sequencing to Identify Causal Genetic Variants Influencing CVD Risk - San Antonio Family Studies | legacy | SAFS | ILLUMINA | R01HL113322 |
| phs000921 | SAGE | Study of African Americans, Asthma, Genes and Environment | 1 | PGX_Asthma | NYGC | 3R01HL117004-02S3 |
| phs000921 | SAGE | ATGC Study of African Americans, Asthma, Genes and Environment | 3 | ATGC | UW NWGC | HHSN268201600032I |
| phs001467 | SAPPHIRE_asthma | Study of Asthma Phenotypes & Pharmacogenomic Interactions by Race-Ethnicity | 3 | ATGC | UW NWGC | HHSN268201600032I |
| phs001207 | Sarcoidosis | Genetics of Sarcoidosis in African Americans | 2 | Sarcoidosis | BAYLOR | 3R01HL113326-04S1 |
| phs001207 | Sarcoidosis | Genetics of Sarcoidosis in African Americans | 3.5 | Sarcoidosis | UW NWGC | HHSN268201600032I |
| phs001446 | SARP | Severe Asthma Research Program | 2 | SARP | NYGC | HHSN268201500016C |
| phs000972 | SAS | Samoan Adiposity Study | 1 | SAS | UW NWGC | HHSN268201100037C |
| phs000972 | SAS | Samoan Adiposity Study | 2 | SAS | NYGC | HHSN268201500016C |
| phs001387 | THRV | Taiwan Study of Hypertension using Rare Variants | 2 | THRV | BAYLOR | 3R01HL111249-04S1, HHSN26820150015C |
| phs001032 | VU_AF | Vanderbilt Genetic Basis of Atrial Fibrillation | 1 | AFGen | BROAD | 3R01HL092577-06S1 |
| phs001237 | WHI | Women's Health Initiative | 2 | WHI | BROAD | HHSN268201500014C |
| NYGC = New York Genome Center; BROAD = Broad Institute of MIT and Harvard; UW NWGC = University of Washington Northwest Genomics Center; ILLUMINA = Illumina Genomic Services; MACROGEN = Macrogen Corp.; BAYLOR = Baylor Human Genome Sequencing Center, WASHU MGI = Washington University McDonnell Genome Institute | | | | | | |

**NHLBI TOPMed: Genetics of Cardiometabolic Health in the Amish (Amish)**

The TOPMed component of the Amish Research Program was supported by NIH grants R01 HL121007, U01 HL072515, and R01 AG18728.

**NHLBI TOPMed: Atherosclerosis Risk in Communities (ARIC)**

The Atherosclerosis Risk in Communities study has been funded in whole or in part with Federal funds from the National Heart, Lung, and Blood Institute, National Institutes of Health, Department of Health and Human Services (contract numbers HHSN268201700001I, HHSN268201700002I, HHSN268201700003I, HHSN268201700004I and HHSN268201700005I).

**NHLBI TOPMed: Barbados Asthma Genetics Study (BAGS)**

Funding for BAGS was provided by National Institutes of Health (NIH) R01HL104608, R01HL087699, and HL104608 S1. The Mount Sinai BioMe Biobank has been supported by The Andrea and Charles Bronfman Philanthropies and in part by Federal funds from the NHLBI and NHGRI (U01HG00638001; U01HG007417; X01HL134588).

**NHLBI TOPMed: Mount Sinai BioMe Biobank (BioMe)**

The Mount Sinai BioMe Biobank has been supported by The Andrea and Charles Bronfman Philanthropies and in part by Federal funds from the NHLBI and NHGRI (U01HG00638001; U01HG007417; X01HL134588). We thank all participants in the Mount Sinai Biobank. We also thank all our recruiters who have assisted and continue to assist in data collection and management and are grateful for the computational resources and staff expertise provided by Scientific Computing at the Icahn School of Medicine at Mount Sinai.

**NHLBI TOPMed: Coronary Artery Risk Development in Young Adults (CARDIA)**

The Coronary Artery Risk Development in Young Adults Study (CARDIA) is conducted and supported by the National Heart, Lung, and Blood Institute (NHLBI) in collaboration with the University of Alabama at Birmingham (HHSN268201300025C & HHSN268201300026C), Northwestern University (HHSN268201300027C), University of Minnesota (HHSN268201300028C), Kaiser Foundation Research Institute (HHSN268201300029C), and Johns Hopkins University School of Medicine (HHSN268200900041C). CARDIA is also partially supported by the Intramural Research Program of the National Institute on Aging (NIA) and an intra-agency agreement between NIA and NHLBI (AG0005).

**NHLBI TOPMed: Cleveland Family Study (CFS)**

Cleveland Family Study is supported by grant numbers-HL 046389; HL113338;1R35HL135818.

**NHLBI TOPMed: Cardiovascular Health Study (CHS)**

Cardiovascular Health Study is supported by contracts HHSN268201200036C, HHSN268200800007C, HHSN268201800001C, N01-HC85079, N01-HC-85080, N01-HC-85081, N01-HC-85082, N01-HC-85083, N01-HC-85084, N01-HC-85085, N01-HC-85086, N01-HC-35129, N01-HC-15103, N01-HC-55222, N01-HC-75150, N01-HC-45133, and N01-HC-85239; grant numbers U01 HL080295, U01 HL130114 and R01 HL059367 from the National Heart, Lung, and Blood Institute, and R01 AG023629 from the National Institute on Aging, with additional contribution from the National Institute of Neurological Disorders and Stroke.

**NHLBI TOPMed: Genetic Epidemiology of COPD Study (COPDGene)**

The COPDGene project described was supported by Award Number U01 HL089897 and Award Number U01 HL089856 from the National Heart, Lung, and Blood Institute. The content is solely the responsibility of the authors and does not necessarily represent the official views of the National Heart, Lung, and Blood Institute or the National Institutes of Health. The COPDGene project is also supported by the COPD Foundation through contributions made to an Industry Advisory Board comprised of AstraZeneca, Boehringer Ingelheim, GlaxoSmithKline, Novartis, Pfizer, Siemens and Sunovion.

**NHLBI TOPMed: Evaluation of COPD Longitudinally to Identify Predictive Surrogate End-points (ECLIPSE)**

The ECLIPSE study (NCT00292552) was sponsored by GlaxoSmithKline.

**NHLBI TOPMed: Boston Early-Onset COPD Study (EOCOPD)**

The Boston Early-Onset COPD Study was supported by R01 HL113264 and U01 HL089856 from the National Heart, Lung, and Blood Institute.

**NHLBI TOPMed: Framingham Heart Study (FHS)**

The Framingham Heart Study (FHS) acknowledges the support of contracts NO1-HC-25195 and HHSN268201500001I from the NHLBI and grant supplement R01 HL092577-06S1 for this research.

**NHLBI TOPMed: ATGC Gene-Environment, Admixture and Latino Asthmatics Study I Asthma (GALAI)**

The Genes-environments and Admixture in Latino Americans (GALA I) Study was supported by the National Heart, Lung, and Blood Institute of the National Institute of Health (NIH) grants R01HL117004 and X01HL134589; study enrollment supported by Sandler Center for Basic Research in Asthma and the Sandler Family Foundation, the American Asthma Foundation, the American Lung Association, the NIH grants K23HL04464 and HL07185, the Resource Centers for Minority Aging Research from the National Institute on Aging, RCMAR P30-AG15272, the National Institute of Nursing Research and the National Center on Minority Health and Health Disparities

**NHLBI TOPMed: Gene-Environment, Admixture and Latino Asthmatics Study (GALAII)**

The Genes-environments and Admixture in Latino Americans (GALA II) Study was supported by the National Heart, Lung, and Blood Institute of the National Institute of Health (NIH) grants R01HL117004 and X01HL134589; study enrollment supported by the Sandler Family Foundation, the American Asthma Foundation, the RWJF Amos Medical Faculty Development Program, Harry Wm. and Diana V. Hind Distinguished Professor in Pharmaceutical Sciences II and the National Institute of Environmental Health Sciences grant R01ES015794 .

The GALA II study collaborators include Shannon Thyne, UCSF; Harold J. Farber, Texas Children's Hospital; Denise Serebrisky, Jacobi Medical Center; Rajesh Kumar, Lurie Children's Hospital of Chicago; Emerita Brigino-Buenaventura, Kaiser Permanente; Michael A. LeNoir, Bay Area Pediatrics; Kelley Meade, UCSF Benioff Children’s Hospital, Oakland; William Rodriguez-Cintron, VA Hospital, Puerto Rico; Pedro C. Avila, Northwestern University; Jose R. Rodriguez-Santana, Centro de Neumologia Pediatrica; Luisa N. Borrell, City University of New York; Adam Davis, UCSF Benioff Children's Hospital, Oakland; Saunak Sen, University of Tennessee and Fred Lurmann, Sonoma Technologies, Inc.

The authors acknowledge the families and patients for their participation and thank the numerous health care providers and community clinics for their support and participation in GALA II. In particular, the authors thank study coordinator Sandra Salazar; the recruiters who obtained the data: Duanny Alva, MD, Gaby Ayala-Rodriguez, Lisa Caine, Elizabeth Castellanos, Jaime Colon, Denise DeJesus, Blanca Lopez, Brenda Lopez, MD, Louis Martos, Vivian Medina, Juana Olivo, Mario Peralta, Esther Pomares, MD, Jihan Quraishi, Johanna Rodriguez, Shahdad Saeedi, Dean Soto, Ana Taveras; and the lab researcher Celeste Eng who processed the biospecimens.

**NHLBI TOPMed: Genetic Studies of Atherosclerosis Risk (GeneSTAR)**

GeneSTAR was supported by grants from the National Institutes of Health/National Heart, Lung, and Blood Institute (U01 HL72518 and HL087698 to LC Becker; HL112064 to RA Mathias) and by a grant from the National Institutes of Health/National Center for Research Resources (M01-RR000052) to the Johns Hopkins General Clinical Research Center.

**NHLBI TOPMed: Genetic Epidemiology Network of Arteriopathy (GENOA)**

Support for GENOA was provided by the National Heart, Lung and Blood Institute (HL054457, HL054464, HL054481, and HL087660) of the National Institutes of Health.

**NHLBI TOPMed: Genetic Epidemiology Network of Salt-Sensitivity (GenSalt)**

The Genetic Epidemiology Network of Salt-Sensitivity (GenSalt) was supported by research grants (U01HL072507, R01HL087263, and R01HL090682) from the NHLBI.

**NHLBI TOPMed: Groningen Genetics of Atrial Fibrillation Study (GGAF)**

The studies that form the GGAF cohort are funded by 5 different sources. The AF RISK study is supported by the Netherlands Heart Foundation (grant NHS2010B233), and the Center for Translational Molecular Medicine. The University Medical Center Groningen supports both the Young-AF and Biomarker-AF studies. The grant 95103007 from ZonMw, the Netherlands Organization for Health Research and Development is funding the GIPS-III trial. The Dutch Kidney Foundation (grant E0.13) and the Netherlands Heart Foundation (grant NHS2010B280) are funding the PREVEND study.

**NHLBI TOPMed: Genetics of Lipid Lowering Drugs and Diet Network (GOLDN)**

GOLDN is supported by U01 HL072524. Whole-genome sequencing in GOLDN was funded by NHLBI grant R01 HL104135 and supplement R01 HL104135-04S1.

**NHLBI TOPMed: Hispanic Community Health Study - Study of Latinos (HCHS_SOL)**

The Hispanic Community Health Study/Study of Latinos is a collaborative study supported by contracts from the National Heart, Lung, and Blood Institute (NHLBI) to the University of North Carolina (HHSN268201300001I / N01-HC-65233), University of Miami (HHSN268201300004I / N01-HC-65234), Albert Einstein College of Medicine (HHSN268201300002I / N01-HC-65235), University of Illinois at Chicago – HHSN268201300003I / N01-HC-65236 Northwestern Univ), and San Diego State University (HHSN268201300005I / N01-HC-65237). The following Institutes/Centers/Offices have contributed to the HCHS/SOL through a transfer of funds to the NHLBI: National Institute on Minority Health and Health Disparities, National Institute on Deafness and Other Communication Disorders, National Institute of Dental and Craniofacial Research, National Institute of Diabetes and Digestive and Kidney Diseases, National Institute of Neurological Disorders and Stroke, NIH Institution-Office of Dietary Supplements.

**NHLBI TOPMed: Heart and Vascular Health Study (HVH)**

The Heart and Vascular Health Study was supported by grants HL068986, HL085251, HL095080, and HL073410 from the National Heart, Lung, and Blood Institute.

**NHLBI TOPMed: Hypertension Genetic Epidemiology Network (HyperGEN)**

 The HyperGEN Study is part of the National Heart, Lung, and Blood Institute (NHLBI) Family Blood Pressure Program; collection of the data represented here was supported by grants U01 HL054472 (MN Lab), U01 HL054473 (DCC), U01 HL054495 (AL FC), and U01 HL054509 (NC FC). The HyperGEN: Genetics of Left Ventricular Hypertrophy Study was supported by NHLBI grant R01 HL055673 with whole-genome sequencing made possible by supplement -18S1.

**NHLBI TOPMed: Jackson Heart Study (JHS)**

 The Jackson Heart Study (JHS) is supported and conducted in collaboration with Jackson State University (HHSN268201300049C and HHSN268201300050C), Tougaloo College (HHSN268201300048C), and the University of Mississippi Medical Center (HHSN268201300046C and HHSN268201300047C) contracts from the National Heart, Lung, and Blood Institute (NHLBI) and the National Institute for Minority Health and Health Disparities (NIMHD).

**NHLBI TOPMed: Mayo Clinic Venous Thromboembolism Study (Mayo_VTE)**

Mayo Clinic Venous Thromboembolism Study is funded by NHLBI grants HL66216 and HL83141 and NHGRI grants HG04735, HG06379, and research support provided by Mayo Foundation.

**NHLBI TOPMed: Multi-Ethnic Study of Atherosclerosis (MESA)**

MESA and the MESA SHARe project are conducted and supported by the National Heart, Lung, and Blood Institute (NHLBI) in collaboration with MESA investigators. Support for MESA is provided by contracts HHSN268201500003I, N01-HC-95159, N01-HC-95160, N01-HC-95161, N01-HC-95162, N01-HC-95163, N01-HC-95164, N01-HC-95165, N01-HC-95166, N01-HC-95167, N01-HC-95168, N01-HC-95169, UL1-TR-000040, UL1-TR-001079, UL1-TR-001420. MESA Family is conducted and supported by the National Heart, Lung, and Blood Institute (NHLBI) in collaboration with MESA investigators. Support is provided by grants and contracts R01HL071051, R01HL071205, R01HL071250, R01HL071251, R01HL071258, R01HL071259, and by the National Center for Research Resources, Grant UL1RR033176. The provision of genotyping data was supported in part by the National Center for Advancing Translational Sciences, CTSI grant UL1TR001881, and the National Institute of Diabetes and Digestive and Kidney Disease Diabetes Research Center (DRC) grant DK063491 to the Southern California Diabetes Endocrinology Research Center.

**NHLBI TOPMed: My Life, Our Future: Genotyping for Progress in Hemophilia (MLOF)**

The My Life, Our Future samples and data are made possible through the partnership of Bloodworks Northwest, the American Thrombosis and Hemostasis Network, the National Hemophilia Foundation, and Bioverativ.

**NHLBI TOPMed: Outcome Modifying Genes in Sickle Cell Disease (OMG-SCD)**

The OMG-SCD study was administrated by Marilyn J. Telen, M.D. and Allison E. Ashley-Koch, Ph.D. from Duke University Medical Center and collection of the data set was supported by grants HL068959 and HL079915 from the NHLBI.

**NHLBI TOPMed: San Antonio Family Study (SAFS)**

Collection of the San Antonio Family Study data was supported in part by National Institutes of Health (NIH) grants R01 HL045522, MH078143, MH078111 and MH083824; and whole genome sequencing of SAFS subjects was supported by U01 DK085524 and R01 HL113323.

**NHLBI TOPMed: SAGE**

The Study of African Americans, Asthma, Genes and Environments (SAGE) was supported by by the National Heart, Lung, and Blood Institute of the National Institute of Health (NIH) grants R01HL117004 and X01HL134589; study enrollment supported by the Sandler Family Foundation, the American Asthma Foundation, the RWJF Amos Medical Faculty Development Program, Harry Wm. and Diana V. Hind Distinguished Professor in Pharmaceutical Sciences II.

The SAGE study collaborators include Harold J. Farber, Texas Children's Hospital; Emerita Brigino-Buenaventura, Kaiser Permanente; Michael A. LeNoir, Bay Area Pediatrics; Kelley Meade, UCSF Benioff Children’s Hospital, Oakland; Luisa N. Borrell, City University of New York; Adam Davis, UCSF Benioff Children’s Hospital, Oakland and Fred Lurmann, Sonoma Technologies, Inc.

The authors acknowledge the families and patients for their participation and thank the numerous health care providers and community clinics for their support and participation in SAGE. In particular, the authors thank study coordinator Sandra Salazar; the recruiters who obtained the data: Lisa Caine, Elizabeth Castellanos, Brenda Lopez, MD, Shahdad Saeedi; and the lab researcher Celeste Eng who processed the biospecimens.

**NHLBI TOPMed: Study of Asthma Phenotypes & Pharmacogenomic Interactions by Race-Ethnicity (SAPPHIRE_asthma)**

The SAPPHIRE study is supported by the Fund for Henry Ford Hospital, the American Asthma Foundation, the National Heart Lung and Blood Institute (R01HL118267, X01HL134589), the National Institute of Allergy and Infectious Diseases (R01AI079139), and the National Institute of Diabetes and Digestive and Kidney Diseases (R01DK113003).

**NHLBI TOPMed: Sarcoidosis**

National Institutes of Health (R01HL113326, P30 GM110766-01)

**NHLBI TOPMed: Samoan Adiposity Study (SAS)**

Samoan Adiposity Study date collection was funded by NIH grant R01-HL093093.

**NHLBI TOPMed: Rare Variants for Hypertension in Taiwan Chinese (THRV)**

The Rare Variants for Hypertension in Taiwan Chinese (THRV) is supported by the National Heart, Lung, and Blood Institute (NHLBI) grant (R01HL111249) and its participation in TOPMed is supported by an NHLBI supplement (R01HL111249-04S1). SAPPHIRe was supported by NHLBI grants (U01HL54527, U01HL54498) and Taiwan funds, and the other cohorts were supported by Taiwan funds.

**NHLBI TOPMed: WHI**

The WHI program is funded by the National Heart, Lung, and Blood Institute, National Institutes of Health, U.S. Department of Health and Human Services through contracts HHSN268201600018C, HHSN268201600001C, HHSN268201600002C, HHSN268201600003C, and HHSN268201600004C.
